# The H4K20 demethylase DPY-21 regulates the dynamics of condensin DC binding

**DOI:** 10.1101/2021.04.11.438056

**Authors:** Laura Breimann, Ana Karina Morao, Jun Kim, David Sebastian Jimenez, Nina Maryn, Krishna Bikkasani, Michael J Carrozza, Sarah E Albritton, Maxwell Kramer, Lena Annika Street, Kustrim Cerimi, Vic-Fabienne Schumann, Ella Bahry, Stephan Preibisch, Andrew Woehler, Sevinç Ercan

**Affiliations:** Department of Biology, Center for Genomics and Systems Biology, New York University, New York, NY, USA; Berlin Institute for Medical Systems Biology, Max Delbrück Center for Molecular Medicine, Berlin, Germany; Institute for Biology, Humboldt University of Berlin, Berlin, Germany; Janelia Research Campus, Howard Hughes Medical Institute, Ashburn, VA, USA

**Keywords:** condensin, transcription, histone modifications, FRAP, Hi-C, *C. elegans*

## Abstract

Condensin is a multi-subunit SMC complex that binds to and compacts chromosomes. Here we addressed the regulation of condensin binding dynamics using *C. elegans* condensin DC, which represses X chromosomes in hermaphrodites for <u>d</u>osage <u>c</u>ompensation. We established fluorescence recovery after photobleaching (FRAP) using the SMC4 homolog DPY-27 and showed that a well-characterized ATPase mutation abolishes its binding. Next, we performed FRAP in the background of several chromatin modifier mutants that cause varying degrees of X-chromosome derepression. The greatest effect was in a null mutant of the H4K20me2 demethylase DPY-21, where the mobile fraction of condensin DC reduced from ∼30% to 10%. In contrast, a catalytic mutant of *dpy-21* did not regulate condensin DC mobility. Hi-C data in the *dpy-21* null mutant showed little change compared to wild type, uncoupling Hi-C measured long-range DNA contacts from transcriptional repression of the X chromosomes. Together, our results indicate that DPY-21 has a non-catalytic role in regulating the dynamics of condensin DC binding, which is important for transcription repression.

## INTRODUCTION

The evolutionarily conserved structural maintenance of chromosomes (SMC) complexes use the energy from ATP hydrolysis to regulate chromosome structure in various nuclear processes (Hirano, 2016). Condensin is an SMC complex that regulates DNA compaction for chromosome segregation during cell division and genome organization for transcription regulation during interphase (Paul et al., 2018a). The current model for how condensins compact DNA is through a process called loop extrusion (Cacciatore and Rowland, 2019; Goloborodko et al., 2016). Unlike a related SMC complex called cohesin, the proteins and chromatin factors that regulate the dynamics of condensin binding are less clear (Paul et al., 2018b). Here we addressed this question using the *Caenorhabditis elegans* dosage compensation system, where an X-specific condensin binding and function is better understood and serves as a model for the metazoan condensins (Albritton and Ercan, 2018).

In *C. elegans*, X chromosome dosage compensation is mediated by a specialized condensin that forms the core of the dosage compensation complex (DCC) (Meyer, 2005). This X-specific condensin (hereafter condensin DC) is distinguished from the canonical condensin I by a single SMC-4 variant called DPY-27 (Csankovszki et al., 2009). The current model of condensin DC binding to the X chromosomes posits that SDC-2, along with SDC-3 and DPY-30, initiate X-specific binding of the complex to a small number of recruitment elements on the X *(rex*) (Albritton et al., 2017; Csankovszki et al., 2004; Jans et al., 2009). Robust binding of condensin DC to the X chromosomes requires multiple *rex* elements (Albritton et al., 2017). The complex binding is enriched at active promoters, enhancers, and other accessible sites (Ercan et al., 2009; Street et al., 2019). Similar to other SMC complexes, condensin DC likely translocates along DNA through loop extrusion and mediates long-range DNA contacts enriched on the X chromosomes (Anderson et al., 2019; Crane et al., 2015; Jimenez et al., 2021). A subset of the strong *rex* sites also serve as blocks to condensin DC movement, insulating DNA contacts and forming loop-anchored topologically associating domains (TADs) (Crane et al., 2015; Jimenez et al., 2021).

Condensin DC physically interacts with DPY-21 (Yonker and Meyer, 2003), a Jumonji domain-containing histone demethylase that converts H4K20me2 to H4K20me1 (Brejc et al., 2017), resulting in increased H4K20me1 and reduced H4K30me2/3 on the X chromosome (Vielle et al., 2012; Wells et al., 2012). This leads to deacetylation of H4K16 mediated by SIR-2.1 (Wells et al., 2012). As a result, the two dosage compensated X chromosomes in hermaphrodites contain higher H4K20me1 and lower H4K16ac levels. Furthermore, condensin DC and *dpy-21* are also required for lower levels of H3K27ac on the X chromosome (Street et al., 2019). An increase of H4K20me1 and decreased acetylation mirror the histone modification changes on metazoan mitotic chromatin (Schmitz et al., 2020), providing a link between canonical condensin and condensin DC binding to chromatin.

In this study, we analyzed the effect of several mutants that regulate H4K20 methylation and H4K16 acetylation on the dynamics of condensin DC binding using fluorescence recovery after photobleaching (FRAP). We established FRAP in *C. elegans* intestine cells using a GFP-tagged DPY-27 and validated the system by demonstrating that condensin DC mobility increases upon depletion of its recruiter SDC-2. We found that introducing a well-characterized mutation in the ATPase domain of DPY-27 eliminated its binding to the X chromosomes as measured by FRAP and ChIP-seq. Mutants that regulate H4K20me and H4K16ac showed subtle effects on condensin DC binding dynamics as measured by FRAP. The most substantial effect was in the *dpy-21* null mutant, which reduced the fraction of mobile DPY-27 from ∼30% to ∼10%. Unlike the null mutant, the *dpy-21 JmjC* catalytic mutant did not affect condensin DC mobility, suggesting that DPY-21 role in regulating condensin DC binding dynamics is non-catalytic. We performed Hi-C analysis in a *dpy-21* null and *(JmjC)* catalytic mutant and observed little change in long-range DNA contacts, including those between the *rex* sites (Brejc et al., 2017). Together, our results suggest that DPY-21 has a noncatalytic role in regulating the dynamics of condensin DC binding to the X chromosomes, which is important for its function in transcription repression.

## RESULTS

### FRAP measurement of condensin DC binding *in vivo*

To analyze condensin DC binding *in vivo*, we used FRAP, which measured functionally relevant dynamics of condensin binding in budding yeast (Thadani et al., 2018) and condensin I and II complexes in human cells (Gerlich et al., 2006; Walther et al., 2018). We set up the FRAP system using DPY-27, the SMC4 homolog that distinguishes condensin DC from I (Fig. 1A). To fluorescently label DPY-27, we added a Halo tag endogenously at the C-terminus using CRISPR/Cas9 genome editing. Unlike *dpy-27* mutants, which are lethal or dumpy, the resulting animals were phenotypically wild-type, indicating that the tagged protein complements protein function. This was also supported by subnuclear localization of DPY-27::Halo, which is typical of X-specific localization of the DCC (Fig. 1B) (Csankovszki et al., 2004; Jans et al., 2009). DPY-27::Halo did not photobleach sufficiently in our hands, and endogenous tagging by GFP did not produce a strong signal. Thus, we turned to expressing a GFP-tagged copy of DPY-27 using a heat-inducible promoter to perform FRAP. First, we characterized the expression of the transgene by incubating adults at 35°C for 1 hour then moving them to the normal growth temperature of 20°C. After 3 hours at 20°C, excess DPY-27::GFP was visible across the nuclei, but after 8 hours, localization was constrained to a subnuclear domain suggesting that the remaining protein bound specifically to the X chromosomes (Fig. 1C).

**Fig. 1).**
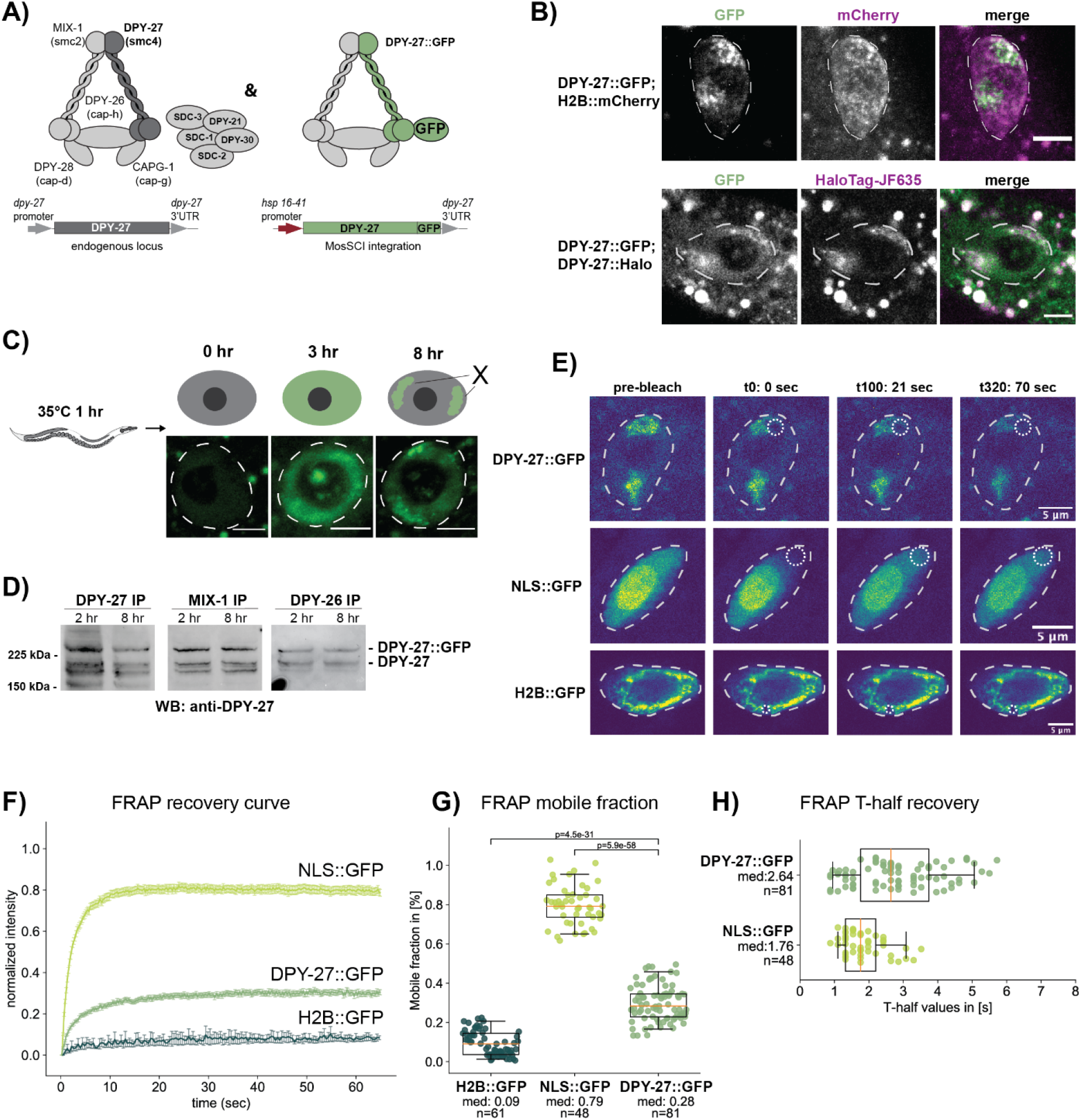
Fluorescence recovery after photobleaching (FRAP) analysis of condensin DC binding. **A)** Left panel illustrates condensin DC along with the rest of the DCC subunits. The right panel indicates the expression of GFP tagged DPY-27 under the control of a heat-shock inducible promoter at the Chr II MosSCI site. **B)** DPY-27::GFP subnuclear localization to the X chromosomes 8 hours after heat-induced expression (top row) was validated by colocalization with the endogenously tagged DPY-27::Halo stained with JF635 HaloTag ligand (bottom row). The scale bars correspond to 5 µm. **C)** Illustration of the heat-shock protocol. Young adult worms were heat-shocked for 1 hour at 35°C, and fluorescence was followed in the large intestinal cells. DPY-27::GFP subnuclear localization is apparent after 8 hours of recovery. Representative example images are shown for each time-point with the nuclear area marked using a white dotted line. The scale bars correspond to 5 µm. **D)** DPY-27::GFP interaction with condensin DC subunits was validated by co-immunoprecipitation with MIX-1 and DPY-26. Young adult worms were used for IP either 2 or 8 hours after heat shock at 35°C for 1 hour and analyzed by western blotting using an anti-DPY-27 antibody. The intensity of the GFP tagged DPY-27 and endogenous protein bands in the DPY-27 IP lane indicates the relative abundance of each protein. The intensity of the GFP tagged DPY-27 and the endogenous protein bands in the other lanes indicates the relative interaction of endogenous and DPY-27::GFP with IPed subunit. **E)** FRAP sequence for intestine nuclei of adult *C. elegans* worms expressing either DPY-27::GFP, NLS::GFP, or H2B::GFP. The first column of images depicts the first image of the pre bleach series (a total of 20 images). The second column shows the first image after the single point bleach with the bleached area indicated by the small dotted circle. The two following columns depict two time points after the bleach point, t100 (21 seconds) and t320 (70 seconds). The scale bars correspond to 5 µm. **F)** Mean FRAP recovery curves from wild-type DPY-27::GFP, H2B::GFP, and NLS::GFP expressing worms. Error bars denote the standard error of the mean (s.e.m.). Number of bleached single intestine nuclei (from at least 3 biological replicates) for each experiment is n = 81 for DPY-27::GFP, n = 48 for NLS::GFP and n= 61 for H2B::GFP. **G)** Mobile fractions for the different GFP tagged proteins or free GFP. The mobile fraction is the lowest for H2B::GFP and the highest for NLS::GFP. The mobile fraction for DPY-27::GFP is ∼28%. P values are from an independent two-sample t-test. **H)** FRAP half-time recovery values for the bleach curves of Fig. 1F. The half-time recovery for NLS::GFP shows a shorter diffusion time than DPY-27::GFP. H2B::GFP is not shown due to the very low recovery of the fluorescence signal during the experimental time frame.

We validated that DPY-27::GFP forms a complex and binds to DNA as expected in three ways. First, we analyzed the localization of DPY-27::GFP after 8 hours of recovery in intestine cells in the presence of the Halo-tagged endogenous protein. DPY-27::GFP colocalized with DPY-27::Halo, indicating proper localization (Fig. 1B, Fig. S1A). Second, DPY-27::GFP immunoprecipitated condensin DC subunits, supporting complex formation (Fig. 1D, Fig. S1B). Third, DPY-27::GFP was enriched on the X chromosomes, and the ChIP-seq binding pattern followed that of DPY-26, the kleisin subunit of condensin DC (Fig. S1C, Fig. 2C,).

**Fig. 2).**
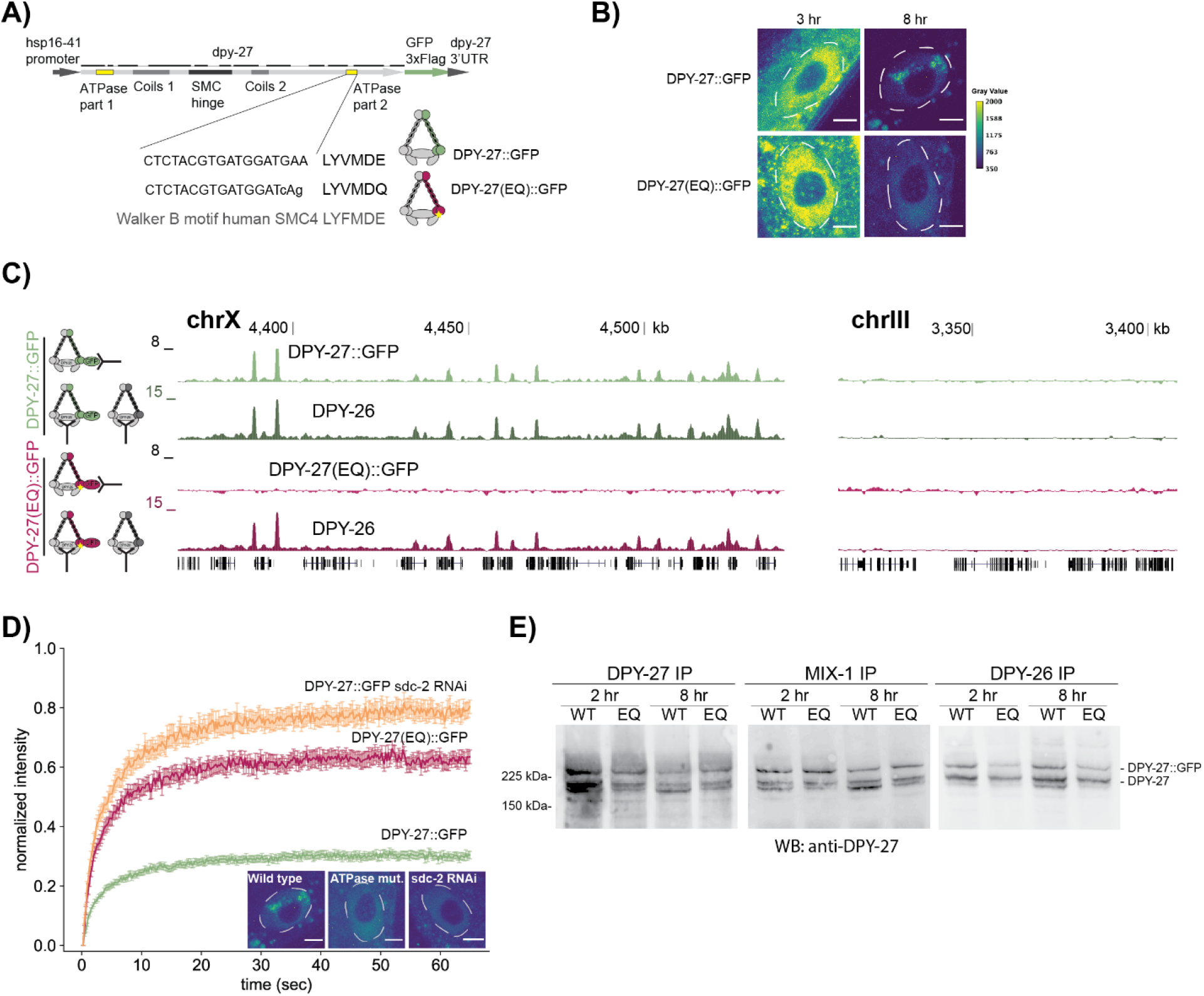
The effect of a conserved SMC ATPase mutation on DPY-27 binding, function, protein stability, and complex formation. **A)** Heat shock inducible GFP tagged *DPY-27(EQ)*. The DNA sequence coding for the conserved Walker B motif and the E to Q mutation are shown below. **B)** Localization of the wild-type and EQ ATPase mutant DPY-27::GFP proteins in intestine cells. Adults were heat-shocked at 35°C for 1 hour and recovered for either 3 or 8 hours. Unlike DPY-27::GFP, ATPase EQ mutant did not show subnuclear localization. The scale bar corresponds to 5 µm. **C)** ChIP-seq analysis of wild-type and ATPase mutant DPY-27(EQ)::GFP using an anti-GFP antibody in embryos. ChIP against DPY-26 was used as a positive control in the same extracts. Unlike the wild-type protein, ATPase mutant failed to bind the X, and both did not localize to the autosomes; a representative region from chromosome III is shown on the right panel. **D)** Mean FRAP recovery curves from DPY-27::GFP, DPY-27(EQ)::GFP and DPY-27::GFP upon SDC-2 RNAi. FRAP was performed ∼8 hr after the heat shock. Error bars denote s.e.m. Number of bleached single intestine nuclei (from at least 3 biological replicates) for each experiment is n = 81 for DPY-27::GFP, n= 37 for DPY-27(EQ)::GFP and n= 32 for DPY-27::GFP sdc-2 RNAi. The small images depict example pictures of intestine nuclei used for FRAP analysis. Unlike DPY-27::GFP, ATPase EQ mutant did not show subnuclear localization, similar to when condensin DC recruiter SDC-2 was knocked down. Scale bars correspond to 5 µm. **E)** Co-immunoprecipitation analysis of condensin DC subunits. Protein extracts were prepared from larvae that were heat-shocked for 1 hour at 35°C and recovered at 20°C for 2 or 8 hours. Immunoprecipitation of condensin DC subunits DPY-27, DPY-26, and MIX-1 was performed, and immunoprecipitated DPY-27::GFP and endogenous protein were analyzed by blotting with an anti-DPY-27 antibody. The intensity of the DPY-27::GFP and endogenous protein bands in the DPY-27 IP lane indicates the relative abundance of each protein. The intensity of DPY-27::GFP and endogenous protein bands in other lanes indicates their relative interaction with each subunit.

We chose intestine cells for performing FRAP, where the nuclei are large due to polyploidy, and subnuclear localization of the complex is easily detected (Fig. 1E). Previous studies also used these cells to analyze condensin DC binding by immunofluorescence (Brejc et al., 2017; Csankovszki et al., 2004; Wells et al., 2012; Yonker and Meyer, 2003). In addition, controlling DPY-27::GFP expression in intestines was easier than in embryos, where nuclei were small (Fig. S1D) and there was variability in heat-induced expression of DPY-27::GFP (Fig. S1E).

To further validate the FRAP assay in the intestine cells, we compared DPY-27::GFP recovery to that of free NLS::GFP and histone H2B::GFP (Fig. 1E-H). FRAP allows two types of quantitative measurements on protein mobility. First is the proportion of mobile molecules, calculated from the percentage of the recovered signal at the bleached area by replacing bleached molecules. Second is the recovery speed, where a fast recovery indicates diffusion, transient binding slows down the recovery, and stable binding increases the immobile fraction (Mueller et al., 2013). As expected, the mobile fraction of free GFP (Fig. 1G) was much higher than that of histone H2B. H2B::GFP minimally recovered during the experiment time frame and was therefore excluded from the half-life recovery plot (Fig. 1H). This result is in line with FRAP experiments in human cell lines reporting a mobile fraction for most H2B-GFP of 4% with a T-half recovery of over 2 hours (Kimura and Cook, 2001).

DPY-27::GFP mobile fraction was ∼30% and half time of recovery ∼2.6 seconds. FRAP results from different experimental set-ups with different imaging settings and analysis strategies can differ significantly (Mazza et al., 2012). However, the time scale for DPY-27::GFP is similar to recovery half-times reported in *Saccharomyces cerevisiae* for Smc4, ∼2 and ∼6 seconds in G1 and M phase respectively (Thadani et al., 2018) and different from residence times reported for human condensin I and II (Gerlich et al., 2006; Walther et al., 2018). During metaphase, human condensin I has a residence time of ∼3 min with a mobile fraction of 80% (Gerlich et al., 2006; Walther et al., 2018). Condensin II, which binds to chromatin throughout the cell cycle, has a residence time of >5 min with a mobile fraction of 40% (Gerlich et al., 2006; Walther et al., 2018). Our results indicate that DPY-27 has a higher chromosome-bound fraction than human condensin I and II but has comparable recovery half-times to those reported in yeast.

### A conserved mutation to the DPY-27 ATPase domain eliminates its binding in the presence of the wild-type protein

If FRAP measures changes in condensin DC binding dynamics, we reasoned that knockdown of its recruiter SDC-2, and a well-characterized ATP hydrolysis mutation that is known to eliminate the function of other SMC4 homologs, should affect DPY-27 binding dynamics. In condensins, the two heads of SMC2 or SMC4 form the two halves of the ATPase domain; each head interacting with the other in the presence of an ATP molecule, hydrolysis of which dissociates the heads (Hirano, 2016). To test if the ATP hydrolysis by DPY-27 is necessary for its binding to DNA, we inserted a walker B mutation (E to Q, Fig. 2A) that nearly eliminates ATP hydrolysis in human (Vian et al., 2018), *Xenopus* (Kinoshita et al., 2015), yeast (Hirano and Hirano, 2004; Thadani et al., 2018), and chicken (Hudson et al., 2008). Unlike wild-type DPY-27::GFP, DPY-27(EQ)::GFP failed to show subnuclear enrichment indicative of localizing to the X chromosome (Fig. 2B, Fig. S2D-E). The conclusion that ATP hydrolysis by DPY-27 is required for its localization to the X was further supported by ChIP-seq analysis of DPY-27::GFP and DPY-27(EQ)::GFP in embryos. Thus, unlike wild-type, the ATPase mutant failed to bind to the X chromosomes in the presence of endogenous DPY-27 (Fig. 2C, Fig. S2A).

Next, we asked if the ATPase mutant improperly interacted with chromatin and showed a dominant-negative effect. The mobility of DPY-27(EQ)::GFP was slightly lower than that of unbound DPY-27::GFP generated by knockdown of condensin DC recruiter SDC-2, thus the mutant may incorrectly associate with chromatin (Fig. 2D, Fig. S2B). Supporting a small dominant-negative effect, mRNA-seq analysis of embryos expressing DPY-27(EQ)::GFP showed slightly higher X chromosome upregulation than those expressing DPY-27::GFP (Fig. S2C). X upregulation upon wild type DPY-27::GFP expression may be due to dosage imbalance within the complex. Additional X upregulation in the EQ mutant may be due to a negative effect on DPY-27 as proposed for SMCL-1, an SMC-like protein with an ATPase hydrolysis mutation (Chao et al., 2017).

To test if the failure of DPY-27(EQ)::GFP to bind is due to its inability to form a complex, we performed co-immunoprecipitation experiments in embryos and young adults (Fig. 2E, Fig. S2F). We noticed that both wild-type and EQ mutant DPY-27::GFP interacted well with MIX-1 (SMC-2 homolog). However, DPY-27::GFP co-IPed better with DPY-26 (kleisin subunit of condensin I and DC) compared to DPY-27(EQ)::GFP, suggesting that ATPase mutation affects SMC-kleisin interaction. Lack of X-specific localization measured by both imaging (Fig. 2B) and ChIP-seq (Fig. 2C) suggests that a combination of inability to form a complex and reduced ATP hydrolysis eliminates binding of DPY-27(EQ)::GFP.

### Recombinant DPY-28 HEAT repeat domain bind to histone H3 and H4 peptides in vitro

We wondered if histone modifications on chromatin regulate dynamics of condensin binding and took a candidate approach, considering histone modifiers that were shown to have a role in *C. elegans* dosage compensation, *set-1* H4K20me1 and *set-4* H4K20me2 transferases, *dpy-21* H4K20me2 demethylase, and *sir-2*.*1* H4K16 deacetylase (Kramer et al., 2015; Wells et al., 2012) (Fig. 3A). A catalytic mutant of DPY-21, *dpy-21(JmjC)* that nearly eliminated its demethylase activity also showed dosage compensation defects, albeit at a lower level than the null mutant (Brejc et al., 2017). Similarly, we found that *sir-2*.*1* null mutant also leads to a slight X derepression (Fig. S3A) and dumpiness, a phenotype indicating dosage compensation problems (Fig. S3B).

**Fig. 3).**
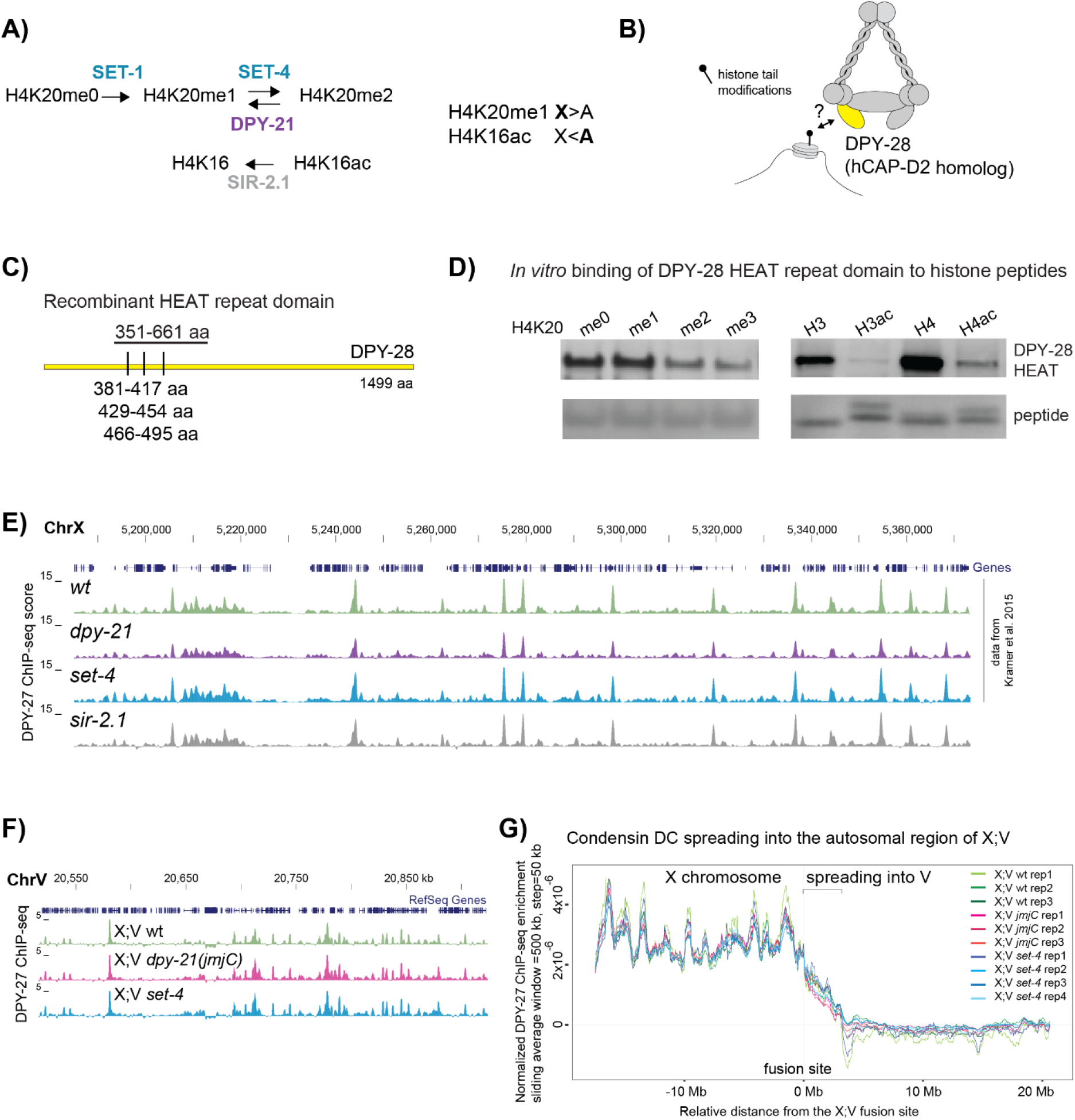
Condensin DC may interact with histone tails, but *set-4, sir-2*.*1*, and catalytic activity of *dpy-21* do not regulate condensin DC binding as measured by ChIP-seq. **A)** Enzymes that regulate H4K20 methylation and H4K16 acetylation. In hermaphrodites, H4K20me1 is increased, and H4K16ac is reduced on the dosage compensated X chromosomes compared to autosomes. *Dpy-21* null is *(e418)* allele with a premature stop codon that eliminates the protein (Yonker and Meyer, 2003), *Dpy-21(JmjC) is the (y607)* allele, a point mutation that nearly abolishes H4K20me2 demethylase activity without eliminating the protein itself (Brejc et al., 2017). *Set-4* null is *(n4600)*, a knockout allele that eliminates H4K20me2/3 (Delaney et al., 2017). *Sir-2*.*1* null is *(ok434)*, a knockout allele that increases H4K16ac (Wells et al., 2012). **B)** Cartoon depicting possible interaction of HEAT repeat-containing domain of DPY-28 (homologous to human hCAPD-2) with histone tail modifications. **C)** Three HEAT repeats annotated by pfam are shown as tick marks. The amino acids 351-661 were purified and used in peptide binding. **D)** In solution peptide binding assay was performed using GST-tagged DPY-28 HEAT domain and biotinylated histone N-terminal tail peptides with indicated modifications. The recombinant protein was incubated with peptides bound to magnetic streptavidin beads, and bound fractions were analyzed using western blot. The streptavidin signal below indicates the amount of peptide in each fraction. **E)** UCSC genome browser shot of a representative region showing similar DPY-27 ChIP-seq patterns in *sir-2*.*1*. Data from wild-type N2, *dpy-21* null, *set-4* null are from (Kramer et al., 2015) and are plotted for comparison. **F)** Genome browser view of DPY-27 ChIP-seq enrichment across the fusion site on the autosomal region of the X;V chromosome in X;V wild-type, *dpy-21(JmjC)* and *set-4* null backgrounds. **G)** A moving average of the DPY-27 ChIP enrichment score is plotted with a window size of 200 kb and step size of 20 kb in X;V fusion strains with wild-type, *dpy-21(JmjC)*, and *set-4* null backgrounds. DPY-27 ChIP-seq data was normalized to reduce variability between replicates by z score standardization ChIP/Input ratios to the background from autosomes I-IV followed by equalization of total ChIP-seq signal to 1 in X:V.

We first considered how condensin DC might interact with histones (Fig. 3B). HEAT repeats, a helical protein structural motif that mediates protein and DNA interactions, are present in the CAPD and CAPG subunits of condensins (Yoshimura and Hirano, 2016). Recombinant HEAT repeat domains from condensin II interacted with H4 peptides monomethylated at lysine 20 (Liu et al., 2010). We asked if the HEAT repeats in condensin I/DC also interact with histone tails. The HEAT repeats in CAPG-1 are predicted to bind DNA (Kschonsak et al., 2017). Thus, we focused on the DPY-28 (CAPD-2 homolog) and identified its HEAT repeat domain using homology to human hCAP-D2 and pfam HEAT predictions (Fig. 3C).

We performed an *in vitro* in-solution peptide binding assay using the recombinant protein (Fig. S3C) and 23 aa N terminal H4 peptides that are unmodified, mono, di, and trimethylated at lysine 20, and unmodified and tetra-acetylated H3 (K4,9,14,18) and H4 (K5,8,12,16) (Fig. S3D). Recombinant DPY-28 HEAT repeat domain interacted with unmodified 23 aa H4 and 20 aa H3 N-terminal peptides (Fig. 3D). Tetra-acetylation and trimethylation of lysine 20 reduced the interaction (Fig. 3D). Thus, histone modifications have the potential to regulate condensin DC interaction with chromatin.

### *SET-4, SIR-2*.*1*, and catalytic activity of DPY-21 do not regulate condensin DC binding

While there is a potential for condensin DC interaction with histones, previous studies showed little effect of chromatin modifier mutants on condensin DC localization, except a slight reduction of DPY-27 ChIP-seq signal across promoters in the *dpy-21* null mutant (Brejc et al., 2017; Kramer et al., 2015; Vielle et al., 2012; Wells et al., 2012). We performed DPY-27 ChIP-seq in *sir-2*.*1* null embryos, and again, did not see a significant difference in condensin DC binding to the X chromosomes compared to wild-type (Fig. 3E). To further rule out the effect of chromatin modifiers, we used X;V fusion chromosomes, where the gradual spreading of condensin DC into the autosomal region may be more sensitive for detecting binding changes (Ercan et al., 2009; Street et al., 2019). We were unable to obtain a homozygous X;V fusion in the *dpy-21* null background, thus we analyzed *dpy-21(JmjC)* and *set-4* null mutants (Fig. 3F, Fig. S3F). In both wild-type, *dpy-21(JmjC)* and *set-4* null backgrounds, ChIP-seq replicates showed variable changes in condensin DC spreading into the autosome (Fig. 3G). Thus, *set-4, sir-2*.*1*, and the catalytic activity of *dpy-21* do not regulate condensin DC binding as measured by ChIP-seq.

### DPY-21 has a non-catalytic activity that increases the mobile fraction of condensin DC

Since the histone modifiers showed little effect on condensin DC binding as measured by ChIP-seq, we used our established FRAP system in mutants and knockdown conditions to study these proteins’ influence on condensin DC dynamics. In *set-1* knockdown, *set-4* null, *sir-2*.*1* null, and *dpy-21(JmjC)* mutants, DPY-27 FRAP recovery was largely similar to that of wild-type, with a small but statistically significant reduction in mobility in *set-4* null (Fig. 4A, Fig. S4A). The most dramatic difference was observed in the *dpy-21* null mutant (Fig. 4A). The *dpy-21* null mutant reduced the percentage of mobile DPY-27::GFP from ∼30% to ∼10% (Fig. 4B). A control experiment bleaching DPY-27::GFP outside of the X indicated that the effect of the *dpy-21* null mutant is largely specific to the X (Fig. S4B). Thus, DPY-21 increases the proportion of mobile condensin DC molecules on the X chromosomes.

**Fig. 4).**
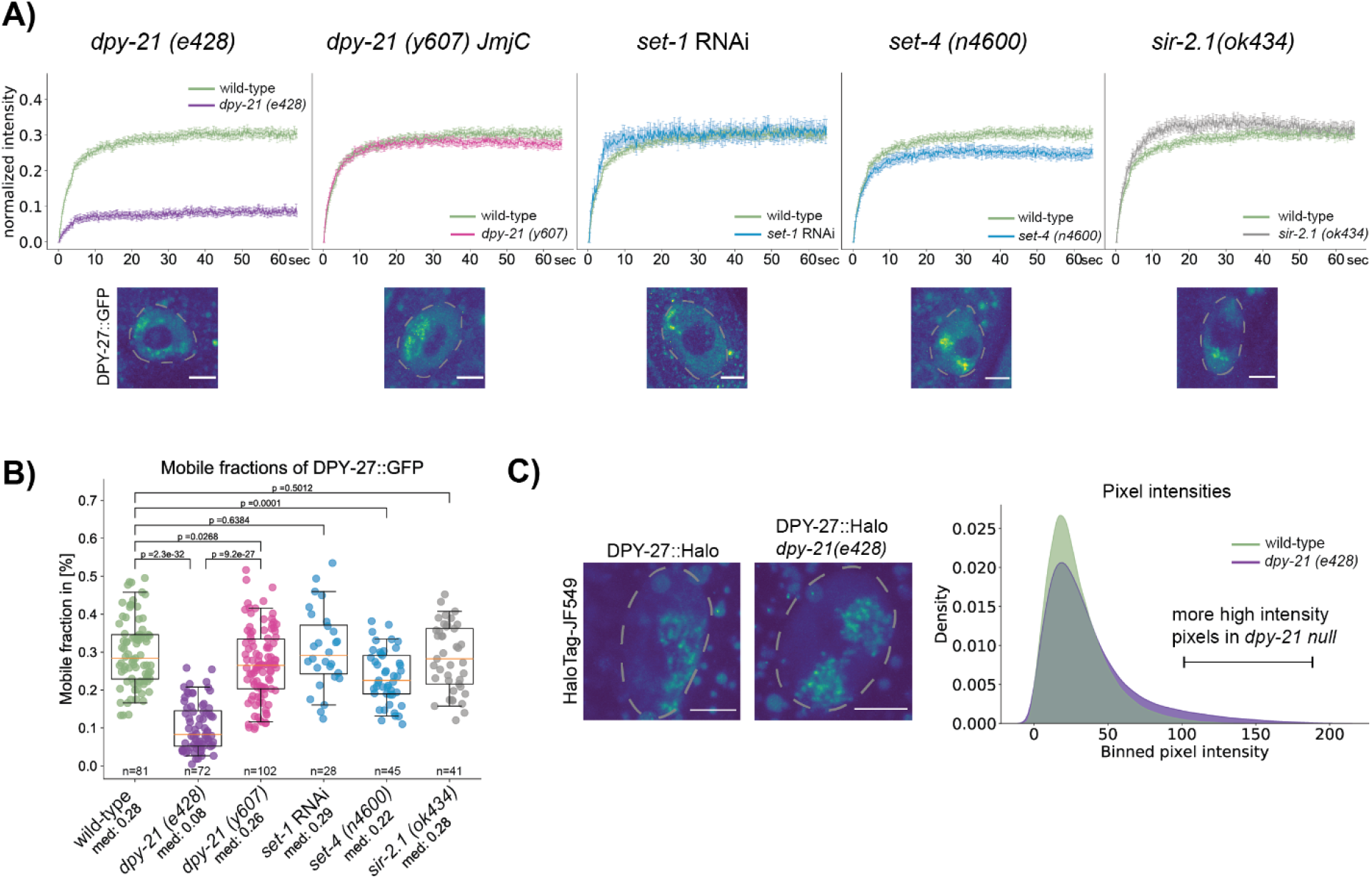
DPY-21 null but not catalytic mutant reduces the proportion of mobile condensin DC. **A)** Mean FRAP recovery curves of DPY-27::GFP in either wild-type (green) or different mutant conditions. Error bars denote s.e.m.. Number of bleached single intestine nuclei (from at least 3 biological replicates) for each experiment is n = 81 for wild-type, n = 72 for *dpy-21 (e428)*, n = 102 for *dpy-21 (y607), set-1 RNAi* n = 28, *set-4 (n4600)* n = 45, *sir-2*.*1 (ok434)* n= 41. Corresponding images of intestine nuclei for each mutant condition are depicted under each FRAP curve. Scale bar = 5 µm. **B)** Mobile fractions calculated from individual replicate FRAP recovery curves in panel A. P values are from an independent two-sample t-test. The number of used images of nuclei is noted under each boxplot. **C)** Analysis of endogenous DPY-27::Halo fluorescent intensity on the X chromosome in wild-type and *dpy-21* null worms. The HaloTag signal of DPY-27 was segmented in 3D and quantified in adult intestine cells in two biological replicates (Fig. S4C). The left panel depicts two example nuclei (marked with a dotted line). Scale bar corresponds to 5 µm. For the wild-type worms, 27 images were analyzed, for the *dpy-21(e428)* mutant images of 35 nuclei were analyzed. The right panel shows the binned mean pixel fluorescence intensity for the two conditions. The distributions of pixel intensities are significantly different in the two conditions according to a Mann-Whitney U test with a p-value of 1.46 *10^−114^.

Previous analysis of condensin DC localization by immunofluorescence in the *dpy-21* null mutant had not reported an effect except an increase in the volume of the X chromosomes in *dpy-21* null and *JmjC* mutants (Brejc et al., 2017; Lau et al., 2014). We wondered if the reduction of mobile condensin DC produces a difference in the confocal imaging of DPY-27::Halo signal compared to wild type. Indeed, we noticed stronger puncta of DPY-27 signal within the X chromosomal domain in the *dpy-21* null mutant, which appears as a long tail of high pixel intensities in the distribution (Fig. 4C, Fig. S4C).

### 3D DNA contacts as measured by Hi-C does not change significantly in the *dpy-21* null

Since *dpy-21* null mutation decreased the number of mobile condensin DC molecules as measured by FRAP, we hypothesized that DPY-21 might act similar to the cohesin unloader WAPL (Haarhuis et al., 2017). To test this idea, we performed Hi-C analysis in *dpy-21* null embryos and repeated Hi-C in *dpy-21(JmjC(y607))* mutants while confirming the strain (Fig. S5D) (Brejc et al., 2017). While a subtle reduction in insulation was observed across a few *rex* sites that act as TAD boundaries, the overall TAD structure was similar to that of wild-type in *dpy-21(JmjC(y607))* and *dpy-21* null embryos on the X chromosomes (Fig. 5A, B) and autosomes (Fig. S5A). The range of DNA interactions in the *dpy-21(JmjC(y607))* and *dpy-21* null mutant are largely similar to the wild-type (Fig. 5C, Fig. S5B) (Brejc et al., 2017).

**Fig. 5).**
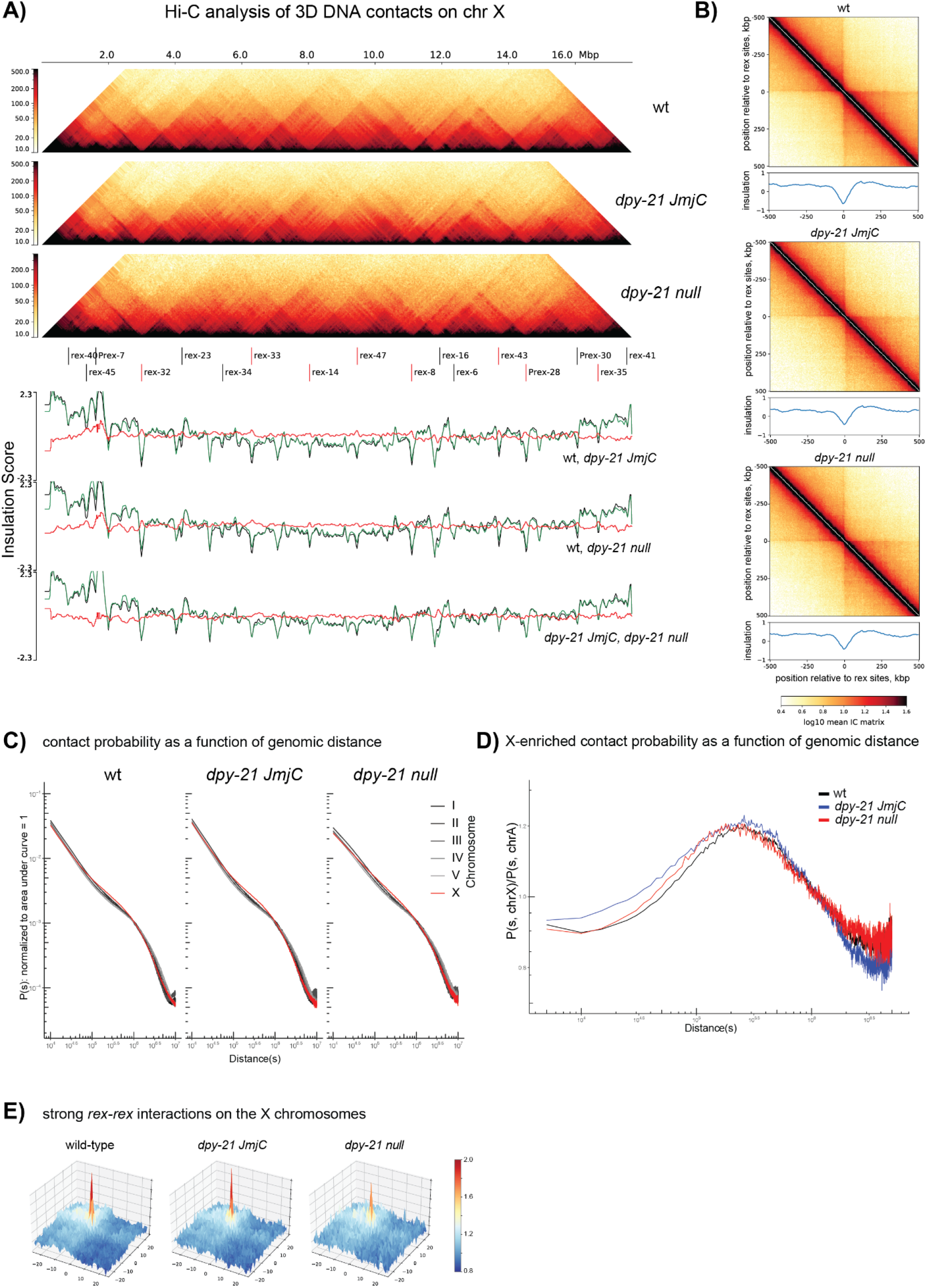
Hi-C analysis of 3D DNA contacts in *dpy-21 JmjC (y607)* and *dpy-21* null mutant embryos. **A)** Hi-C heatmap and insulation scores of chromosome-X showing wild-type, the *dpy-21 JmjC*, and the *dpy-21* null mutant. The 17 strong *rex* sites are annotated in (Albritton et al., 2017), 8 of which were annotated as DCC-dependent boundary *rex* sites (red) in (Anderson et al., 2019). The insulation scores and their subtractions for three possible pairwise comparisons are shown below. **B)** Pile-up analysis showing the average Hi-C map and the insulation scores +/- 500-kb surrounding the annotated 17 strong *rex* sites. **C)** Distance decay curve showing the relationship between 5-kb binned genomic separation, s, and average contact probability, P(s) computed per chromosome. **D)** X-enriched chromosomal contacts are visualized by an X-A normalized distance decay curve. For every genomic separation s, the unity normalized contact probability of X-chromosome, P(s,chrX), is divided by that of autosomes, P(s,chrA). **E)** Meta-’dot’ plot showing the average strength of interactions between pairs of *rex* sites on a distance-normalized matrix. For 17 strong *rex* sites, a total of 33 *rex-rex* pairs located within 3 Mb of each other were used.

To highlight condensin DC-mediated X-specific 3D contacts, we normalized contact frequency across the same distance on the X to autosomes. This analysis reaffirmed that compared to autosomes, DNA contacts between 50 kb to 1 Mb range (approximated based on X/A > 1) are more frequent on the X (Fig. 5D, Fig. S5C). We reasoned that if DPY-21 protein functions as the unloader for condensin DC, a rightward shift in X-enriched contacts would be observed in *dpy-21* null as condensin DC stays loaded on DNA to form larger loops. However, the enriched contacts remained largely unaffected in both *JmjC* and null mutant (Fig. 5D). Furthermore, in contrast to stronger loops observed in the cohesin unloader WAPL mutant, interactions between *rex* sites weakened in the *dpy-21* null mutant (Fig. 5E). Thus, we conclude that DPY-21 does not act as a condensin DC unloader.

## DISCUSSION

*In vivo* and *in vitro* studies show that SMC complex function requires ATPase activity (Hassler et al., 2018; Hirano, 2016). In *C. elegans* condensin DC, four out of five subunits are also within the condensin I complex, thus their functional homology is apparent (Csankovszki et al., 2009). The single subunit that distinguishes condensin DC from condensin I is DPY-27, the SMC4 homolog (Csankovszki et al., 2009; Hagstrom et al., 2002). Here we showed that a single amino acid mutation that has been shown to slow down ATP hydrolysis and impair the function of SMC4 proteins in other organisms also eliminates DPY-27 binding to the X chromosomes (Fig. 2). This observation adds to evidence that the evolutionarily conserved SMC complex activity is conserved in condensin DC (Albritton and Ercan, 2018; Lau and Csankovszki, 2014; Wood et al., 2010).

Although ATPase activity is strictly conserved, there may be differences in how different SMC complexes and organisms are affected by ATPase mutations. In *Xenopus* extracts, incorporating the EQ mutation in SMC-2 and SMC-4 did not abolish loading to chromosomes analyzed by immunofluorescence (IF) (Kinoshita et al., 2015). Similar results were obtained in chicken cell culture and yeast where SMC-2 and SMC-4 EQ single mutants were able to bind chromosomes at levels comparable to the WT but were not competent in chromosome compaction (Hudson et al., 2008; Thadani et al., 2018). In *Bacillus subtilis*, ChIP-seq experiments showed that the EQ mutant SMC bound to *parS* loading sites but had reduced spreading along the chromosome (Minnen et al., 2016). Similarly, mammalian EQ mutant cohesin binding at loading sites was less affected than at CTCF sites (Vian et al., 2018). Thus, different binding modes may have different ATPase requirements, and although the EQ mutation reduced ATP hydrolysis in all SMC complexes analyzed so far, future work is needed to characterize the specific effect of this mutation on condensin DC.

In addition to DNA loop extrusion, ATPase activity may also contribute to SMC complex formation and stability *in vivo*, perhaps by controlling the structural changes that occur through the cycle of ATP binding and hydrolysis (Lee et al., 2020). While in chicken, no measurable effect of ATPase mutation was reported for complex formation measured by pull-down experiments (Hudson et al., 2008), in budding yeast, ATP binding mutation reduced the interaction between SMC-4 and the kleisin subunit (Thadani et al., 2018), and in *B. subtilis*, ATPase mutations reduced the SMC homodimer’s proper interaction with the ScpA bridging protein as measured by crosslinking assay (Wilhelm et al., 2015). We have also noticed reduced co-IP interaction with the kleisin subunit by DPY-27(EQ). These observations suggest that the ATPase cycle affects the formation of condensins *in vivo*.

### Enrichment and depletion of H4K20me1 and H4K16ac on the X chromosomes have little effect on condensin DC binding in vivo

*In vitro*, condensin prefers binding to free DNA *(Kong et al., 2020; Kschonsak et al., 2017; Piazza et al., 2014)*, and *in vivo* ChIP-seq analysis of condensins in various organisms revealed that condensins are accumulated at accessible regions of the genome (Jeppsson et al., 2014; Uhlmann, 2016). Interestingly, a recent study found that condensin is able to extrude DNA fragments containing 3-4 nucleosomes, and the nucleosomes increased the velocity and processivity of condensin II *in vitro* (Kong et al., 2020). Thus, nucleosomes may regulate the ATPase-dependent movement of condensin. In addition to nucleosomes themselves, chromatin modifications, histone variants, and linker histone were proposed to regulate condensin binding (Choppakatla et al., 2021; Kim et al., 2009; Kimura and Hirano, 2000; Liu et al., 2010; Petty et al., 2009; Tada et al., 2011; Tanaka et al., 2012; Yuen et al., 2017).

The potential for HEAT repeat domains in CAP-D3 and CAP-G2 to interact with histones was put forward for human condensin II (Liu et al., 2010). Here, we found that the recombinant HEAT repeat domain of DPY-28 interacts with histone H3 and H4 tail peptides (Fig. 3). Yet, mutants that reduce X-enrichment of H4K20me1 and increase X-depletion of H4K16ac did not affect condensin DC binding as measured by ChIP-seq and showed subtle changes in FRAP (Figs 3, 4). Thus, enrichment and depletion of H4K20me1 and H4K16ac on the X do not control condensin DC binding. It remains unclear if the combined effects of multiple histone modifications, variants, and linker histones on the X chromosomes regulate condensin DC binding.

### A non-catalytic activity of DPY-21 regulates the dynamics of condensin DC binding and is required for transcription repression on the X chromosomes

DPY-21 is an H4K20me2 demethylase that interacts with condensin DC and is important for dosage compensation (Brejc et al., 2017; Yonker and Meyer, 2003). Comparison of the null and catalytic mutants indicated that DPY-21 plays both a structural and catalytic role in X chromosome repression (Brejc et al., 2017). The catalytic role of *dpy-21* decreases H4K20me2/3 and increases H4K20me1 on the X chromosomes and contributes to repression. Here, we showed that DPY-21’s non-catalytic role increases the mobile fraction of condensin DC on the X chromosomes, which is critical for transcription repression.

How do the catalytic and noncatalytic activities of DPY-21 contribute to repression? DPY-21 mediated enrichment of H4K20me1 leads to reduction of H4K16ac on the X chromosomes, which may reduce binding of general activator(s), contributing a portion of the observed 2-fold repression provided by condensin DC (Sheikh et al., 2019). Our work suggests that a non-catalytic activity of DPY-21 contributes to repression by regulating the kinetics of condensin DC diffusion. In the *dpy-21* null mutant, but not in the *JmjC* mutant, the fraction of mobile condensin DC reduced from ∼30% to ∼10%. Interestingly, in the *dpy-21* null mutant, condensin DC binding to promoters slightly decreases (Kramer et al., 2015), and the DPY-27::Halo signal shows higher intensity spots. It is possible that, without DPY-21, condensin DC is more frequently “trapped” in an immobile configuration that reduces condensin DC presence and activity at promoters that represses transcription initiation.

How does DPY-21 increase the proportion of the mobile condensin DC complexes? Hi-C analysis in the *dpy-21* null mutant argues against a role akin to the cohesin unloader WAPL (Haarhuis et al., 2017; Nuebler et al., 2018). A previous study reported that in the *JmjC* catalytic mutant, long-range interactions between *rex* sites were diminished, and shorter-range interactions increased (Brejc et al., 2017). Qualitatively, we replicated the enrichment of shorter-range interaction (Fig. 5D) in the *JmjC* catalytic mutant. However, the observed effect was small, and there was no significant change in long-range interactions between *rex* sites (Fig. 5E). The Hi-C measurements in the *dpy-21* null mutant differed little from the wild type, suggesting that DPY-21 protein does not regulate condensin DC-mediated long-range 3D DNA contacts on the X.

The noncatalytic activity of DPY-21 may be structural, similar to those reported for other histone-modifying enzymes, including demethylases. For example, a range of noncatalytic activities for the Lysine-specific demethylase 1 (LSD1 or KDM1A) have been discovered, including the role of LSD1 as a scaffolding protein, destabilizing other proteins by promoting self-ubiquitylation, inhibiting autophagy, or protecting other proteins from proteasome-dependent degradation (Gu et al., 2020; Miller et al., 2020). JmjC domain-containing demethylases also show noncatalytic activities. Kdm2b, the H3K36 demethylase, recruits PRC1 to unmethylated CpG islands via its zinc finger domain (He et al., 2013). Similarly, the H3K36 demethylase dKDM4A in *Drosophila* regulates heterochromatin position-effect variegation independent of its catalytic activity (Colmenares et al., 2017). In fission yeast, overexpression of the histone demethylase Epe1 causes heterochromatin defects by recruiting the histone acetyltransferase complex SAGA, independent of the demethylase activity (Bao and Jia, 2019). For DPY-21 so far, structural work is limited to 407 aa that includes the JmjC domain (Brejc et al., 2017). Secondary structure prediction tools suggest that the rest of the 1641 aa long protein is highly unstructured. Intrinsically disordered protein domains promote protein-protein and protein-nucleic acid interactions (Davey, 2019). DPY-21 could directly or indirectly interact with condensin DC, perhaps through regulating its binding to histone tails and controlling its mobility.

Interestingly, while X chromosomes are upregulated ∼2-fold in the *dpy-21* null mutant, Hi-C showed minimal change at the chromosome-wide level. This could be due to condensin-DC mediated DNA loops not being sufficient for repression or the lack of temporal or gene-level resolution of Hi-C data. Higher-resolution assays such as Micro-C may detect shorter-range DNA contacts that may be relevant to condensin-mediated repression (Swygert et al., 2019). Alternatively, the temporal dynamics of condensin DC may be important for repression, which could be addressed by high-resolution live imaging of condensin DC association with DNA. Here, our results suggest that the dynamics of condensin DC binding to chromatin is important for its function, and DPY-21 regulates both histone modifications and condensin DC mobility to repress X chromosome transcription (Fig. 6).

**Fig. 6).**
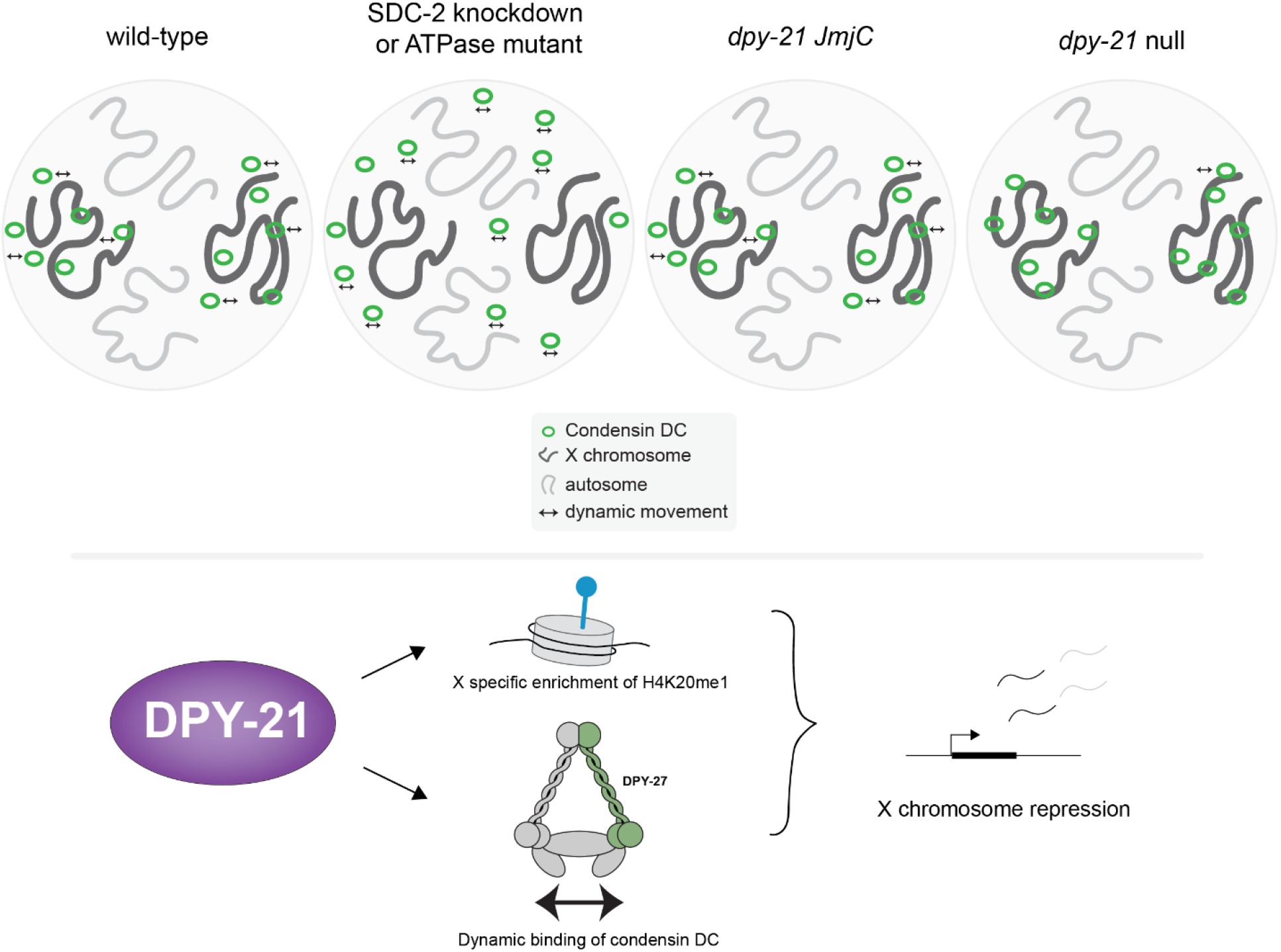
Summary of results and DPY-21 function in condensin DC-mediated X chromosome repression. In a wild-type hermaphrodite cell, condensin DC binds dynamically and specifically to the X chromosomes. This binding is disrupted by a knockdown of its recruiter SDC-2 or a single amino acid mutation in the ATPase domain of DPY-27. Condensin DC may interact with histone tails through HEAT-repeats within DPY-28. H4K20me2 demethylase, DPY-21 has a dual function in X chromosome repression. The catalytic activity reduces H4K20me2/3 and increases H4K20me1 on the X. This leads to reduced H4K16ac and contributes to repression. Both catalytic and non-catalytic activities of DPY-21 do not regulate the X-enriched long-range 3D contacts measured by Hi-C. The non-catalytic activity of DPY-21 increases the mobility of condensin DC molecules, which is important for transcription repression. In the *dpy-21* null condition, both catalytic and non-catalytic activities are eliminated, resulting in stronger X chromosome derepression.

## MATERIALS AND METHODS

### Strains and Worm Growth

A list of strains, genotypes, and primer sequences are provided in Tables S1, S2. Unless noted, worms were grown and maintained using standard methods at 20-22°C on NGM plates containing OP50-1 strain of *E. coli* as food.

#### Generation of DPY-27::GFP and DPY-27(EQ)::GFP strains

An inducible GFP-tagged copy of DPY-27 was expressed from the chrII MosSCI site (∼8.4 Mb) (Frøkjær-Jensen et al., 2008) under the control of a heat-shock inducible *hsp 16-41* promoter and the *dpy-27* 3’ UTR. The *Hsp 16-41* promoter was amplified from pCM1.57 using primers SE123 F&R, and *dpy-27* 3’UTR was amplified from genomic DNA using primers SE124F&R and were inserted into pCFJ151 at the XhoI site. The resulting plasmid contained a SphI site between the promoter and the 3’ UTR, which was used for NEB Infusion cloning with the full-length *dpy-27* and a GFP-3xflag sequence. Amplification of the *dpy-27* sequence was done from genomic DNA using primers SE135F&R. GFP-3x flag sequence was amplified from a plasmid kindly provided by Susan Strome, using primers SE136 F&R. ATPase mutagenesis of DPY-27 was performed by incorporating the E to Q mutation at the conserved ATPase domain as shown in Fig. 2A.

#### Generation of DPY-27::Halo strain

The CRISPR/Cas9 system was used to insert the Halo tag at the C-terminus of DPY-27 (Dokshin et al., 2018). A 20 bp crRNA (LS37) was designed to target the end of the last *dpy-27* exon. The dsDNA donors consisting of a 15 bp flexible linker (GlyGlyGlyGlySer) and the Halo tag flanked by 35 bp homology arms were generated by PCR using 5’ SP9 (TEG) modified primers AM29F&R and pLS19 as a template. The injection mix containing *S*.*pyogenes* Cas9 3NLS (10 μg/μl, IDT), crRNA (2 nmol, IDT), tracrRNA (5 nmol, IDT), dsDNA donors, and pCFJ90 (pharynx mCherry marker) was prepared as previously described (Dokshin et al., 2018). ∼40 F1s that were positive for the co-injection marker were transferred to individual plates and allowed to have progeny. F2 progeny was screened by PCR with primers LS40F&R. Sanger sequencing of positive PCR products showed in-frame insertion of the Halo tag along with 18 bp of unknown sequence that did not affect the function of the tagged protein (Table S5).

#### Generation of X;V, set-4(n4600) and X;V,dpy21(y607 JmjC) strains

ERC57 (*set-4 (n4600) II; X;V (ypT47)*) strain was generated by crossing YPT47 with the set-4 null deletion mutant strain MT14911. For X;V, *dpy21(*same as *y607 JmjC)* strain, a single amino acid substitution (H1452A), that disrupts the demethylase activity of *dpy-21* (Brejc et al., 2017), was incorporated in the *X;V (ypT47)* strain using CRISPR/Cas9. A 200 bp single-stranded oligonucleotide repair template (BR16_oligo) was used to change the codon 1452 from CAC to GCC. The introduction of the changed codon generated a NotI restriction site that was used to screen and confirmed by Sanger sequencing (Table S6). BR17F&R primers amplify a 514 bp region that encompasses the mutation site, and NotI digestion generates two fragments of 216 bp and 298 bp only in the mutated allele.

### Genomic Data Access

The new genomic data is available at Gene Expression Omnibus (GEO) series numbers GSE169458, and individual accession numbers of the new and published data sets used in this study are listed in Tables S4, S7, and S8.

### ChIP-seq

For the ChIP-seq analyses of GFP tagged DPY-27 in embryos, gravid adults were heat-shocked at 35°C for 30 min and transferred to room temperature for two hours for recovery. Embryos were collected by bleaching, and ChIP was performed as described previously (Ercan et al., 2007). Two micrograms of anti-GFP (Abcam ab290) and anti-DPY-26 antibodies were used with 1-2 mg of embryo extract. Detailed antibody information is given in Tables S3and S7. The ChIP-seq analysis of the X;V fusion strains was performed in early L3 larvae by hatching embryos in M9 overnight. The next day, L1s were plated on NGM media containing HB101 bacteria and incubated at 20°C for ∼24 hours. ChIP in larvae was performed by grinding frozen larvae a few minutes in mortar and pestle cooled in liquid nitrogen, followed by crosslinking in PBS containing 1% formaldehyde for 10 min, quenching with 125 mM glycine for 5 min, and preparing ChIP extract as in embryos. X;V wt rep2 was prepared by live crosslinking larvae. Two micrograms of anti-DPY-27 were used with 1-2 mg of extract per ChIP. Half of the ChIP DNA and approximately 20-80 ng of the input control DNA were used to make Illumina TruSeq libraries as previously described (Albritton et al., 2017). For each data set, at least two biological replicates were generated, as listed in Table S7. Single-end sequencing was performed in Illumina HiSeq500 or NextSeq.

ChIP-seq data analysis: We used bowtie2 version 2.3.2 to align 50-75 bp single-end reads to WS220 with default parameters (Langmead and Salzberg, 2012). Bam sorting and indexing was performed using samtools version 2.1.1 (Ramirez-Gonzalez et al., 2012). BamCompare tool in Deeptools version 3.3.1 was used to normalize for the sequencing depth using CPM and create ChIP/Input ratios with a bin size of 10 bp and 200 bp read extension (Ramírez et al., 2016). Only reads with a minimum mapping quality of 20 were used, and mitochondrial DNA, PCR duplicates, and blacklisted regions were removed (Ho et al., 2014). The average coverage data was generated by averaging ChIP-Input enrichment scores per 10 bp bins across the genome. For alignments and sliding window analysis of replicates, ChIP/Input ratios were z-scored using the standard deviation and mean of autosomes or chromosomes I to IV in normal and X;V karyotypes, respectively.

### mRNA-seq

mRNA-seq analysis of *sir-2*.*1* null mutant strain VC199 (*sir-2*.*1*) was performed as described and compared to previously published mRNA-seq data (Kramer et al., 2015). Briefly, embryos and L2/L3 larvae were collected for at least three biological replicates. After collection, worms were stored in Trizol (Invitrogen). RNA was purified using the manufacturer’s protocol after freeze-cracking samples five times. RNA was cleaned up using Qiagen RNeasy kit, and mRNA was purified using Sera-Mag Oligo (dT) beads (Thermo Scientific) from 1 µg of total RNA. Stranded Illumina libraries were prepared as described (Kramer et al., 2015), and sequencing was done with Illumina HiSeq-2000 to produce single-end 50-75 bp reads. We aligned reads to the WS220 genome version using Tophat version 2.1.1 with default parameters (Kim et al., 2013). Count data was calculated using HTSeq version 0.6.1 (Anders et al., 2015) and normalized using the R package DESeq2 (Love et al., 2014).

### Hi-C

CB428 (*dpy-21(e428))* and TY5686 (*dpy-21(y607*)) gravid adults were bleached to isolate embryos, which were crosslinked in 50 mL M9 containing 2% formaldehyde, washed with M9 and PBS, and pelleted at 2000 g 1 min to store at -80°C. Approximately 50 µl of the embryo pellet was resuspended and crosslinked a second time using the same conditions, washed once with 50 mL 100mM Tris-Cl pH 7.5 and twice with 50 mL M9. The embryo pellet was resuspended in 1 ml embryo buffer (110 mM NaCl, 40 mM KCl, 2 mM CaCl2, 2 mM MgCl2, 25 mM HEPES-KOH pH 7.5) containing 1 unit chitinase (Sigma) and digested approximately 15 minutes. Blastomeres were then washed with embryo buffer twice by spinning at 1000g 5 min. The pellet was resuspended in 1 mL Nuclei Buffer A (15 mM Tris–HCl pH 7.5, 2 mM MgCl2, 0.34 M Sucrose, 0.15 mM Spermine, 0.5 mM Spermidine, 1 mM DTT, 0.5 mM PMSF, 1xCalbiochem Protease Inhibitor cocktail I, 0.25% NP-40, 0.1% Triton X-100), centrifuged at 1000 g for 5 minutes at 4°C then resuspended in 1.5 mL Nuclei Buffer A. The embryos were dounced ten times with a loose pestle A and ten times with a tight pestle B. The cellular debris was spun down 1 min at 200 g. The supernatant containing nuclei was kept on ice. The pellet was resuspended in 1.5 mL Nuclei Buffer A, and the douncing process was repeated four times. Each supernatant was checked for absence of debris by DAPI stain and pooled and spun down at 1000 g for 10 mins at 4°C. Approximately ∼20 µl of nuclei were used to proceed to the Arima Hi-C kit, which uses two 4-base cutters, DpnII (^GATC) and HinfI (G^ANTC), followed by KAPA Hyper Prep Kit for library preparation per the protocol provided by Arima. Paired-end Illumina sequencing was performed with Nextseq or Novaseq.

Hi-C data analysis: 150 bp reads were trimmed using fastx toolkit version 0.0.14 to match replicates generated by 100-bp paired-end sequencing. The Hi-C data was mapped to ce10 (WS220) reference genome using default parameters of the Juicer pipeline version 1.5.7 (Durand et al., 2016). Because Hi-C data generated from the Arima Hi-C kit used two restriction enzymes, dpnII (^GATC) and hinfI (G^ANTC), while the published Hi-C data used only one, dpnII (^GATC), the corresponding restriction sites files were used for the juicer pipeline. The mapping statistics from the inter_30.txt output file are provided in Table S8. The inter_30.hic outputs were converted to h5 using the hicConvertFormat of HiCExplorer version=3.5.1 for genome-wide normalization and sample-to-sample depth normalization. (Ramírez et al., 2018; Wolff et al., 2018; Wolff et al., 2020). The inter_30.hic files were first converted to cool files, and the correction method was removed using the --correction_name none option. Then, cool files were converted to h5 files to be used in HiCExplorer. The replicates of the same experimental condition were combined using hicSumMatrices. The count values of each replicate were normalized to match those of the most shallow matrix using hicNormalize with the option --smallest. The same method was used for the summed matrices. Lastly, the hicCorrectMatrix function was applied to each matrix to correct for sequencing bias with the following parameters: --correction_method ICE, -t 1.7 5, --skipDiagonal, --chromosomes I II III IV V X. The distance decay curves were generated by computing the average contact for a given distance using the 5000 bp-binned normalized matrix using hicPlotDistVsCounts with parameters --perchr, maxdepth 20,000,000. The outputs from --outFileData were plotted in R. The curves were normalized to unity to compare different samples by setting the sum of contacts in the distance range of 5000 bp to 4 Mb range to 1 for each chromosome. To analyze X-specific changes, we calculated P(s,chrX)/P(s,chrA) by dividing the P(s) of the X chromosome by the average P(s) of all autosomes at every distance, s. The insulation scores were computed using the 10kb-binned normalized matrix with the function hicFindTADs using parameters: --correctForMultipleTesting fdr, --minDepth 80000, --maxDepth 200000, --step 40000. The meta-loops were computed using the 10 kb-binned normalized matrix with the hicAggregateContacts function of hicexplorer with parameters: --range 100000:3000000, --avgType mean, --transform obs/exp, --plotType 3d, --vMin 0.8 --vMax 2 --BED 17 strong *rex*es (Albritton et al., 2017). A 400 bp window for the 17 strong rex sites defined in (Albritton et al., 2017) was used as center regions with an additional 250 kb up and downstream regions. The pileup analysis at *rex* sites was done using cooltools (https://github.com/open2c/cooltools) by converting the corrected matrix from hicexplorer format to cool format using hicConvertFormat function.

### Immunoprecipitation and Western blots

Immunoprecipitations (IPs) of GFP-tagged DPY-27 proteins were performed from protein extracts prepared using 200 µL of young adult worms heat-shocked at 35°C for one hour and let to recover at 20°C for the indicated times. For IPs from embryos, heat-shocked adults were bleached after recovery to obtain ∼100 µl embryos. Worms were dounced in lysis buffer (40 mM HEPES pH 7.5, 10% glycerol, 150 mM NaCl, 1 mM EDTA, and 0.5% NP-40) complemented with protease inhibitors (Calbiochem cocktail I) and sonicated for 5 min (30 sec on and 30 sec off in a Bioruptor). Extracts were centrifuged at 17,000 g for 15 min at 4°C, and 2 mg of protein were incubated overnight with 2-3 µg of the indicated antibody. Immunocomplexes were collected with protein A Sepharose beads at 4°C for 2 hours. Beads were washed thrice with 1 ml of immunoprecipitation buffer (50 mM HEPES-KOH pH 7.6, 1 mM EDTA, 0.05% Igepal, and 150 mM NaCl). IPed proteins were eluted by boiling in SDS sample buffer and analyzed by SDS-PAGE and immunoblotting using an anti-DPY-27 antibody (1:2000). Detection was performed using ECL Plus reagents (#PI80196, ThermoFisher).

### Worm size analysis

Quantification of the worm size was performed in the young adult stage. Worms were allowed to lay eggs for 4 hours, and the progeny was grown at 20°C to a young adult stage. Worms were washed with M9, anesthetized with 10 mM levamisole, and placed on a fresh NGM plate without OP50 to achieve an even and clear background. Worms were singled with an eyelash, and images of about 30 worms were acquired using a Dino-Lite eyepiece camera (AM7025X) on a Zeiss stereomicroscope with a 1X magnification. For analysis, the background was subtracted using Fiji (Schindelin et al., 2012) with a rolling ball radius of 50 px (light background). The Fiji plugin WormSizer (Moore et al., 2013) was used to analyze the worms’ size and width, and plots were created using Python. (https://github.com/ercanlab/2021_Breimann_et_al)

### RNAi conditions

For RNAi experiments, bacteria strains from the Ahringer RNAi library were verified by Sanger sequencing and used for knockdown experiments (*set-1, sdc-2*, as well as *pop-1* (controls for efficiency of the RNA plates) and empty vector (negative control). Single colonies of bacteria were picked and grown in 10 mL LB with 50 μg/mL ampicillin overnight (at 37°C shaking at 300 rpm), then transferred to a 400 mL LB with 50 μg/mL ampicillin culture and after 2 hours when the culture reached OD ∼1 induced with 0.1 mM ITPG and grown for another 3 hours. Bacteria were concentrated 10-fold and seeded onto 10 cm NGM plates supplemented with 50 μg/mL ampicillin, 2 μg/mL tetracycline, and 1mM IPTG. Worms were synchronized by bleaching, and L1s were placed on the seeded plates. Worms were used for FRAP experiments after 72 hours at 20°C (young adult stage). FRAP experiments for the *set-1* RNAi condition were performed in germline-less worms, indicating successful protein knockdown (Vielle et al., 2012).

### Heat shock, fluorescent labeling, and mounting worms for imaging

JF549-HaloTag and JF635-HaloTag ligands were a generous gift from Luke D. Lavis and Jonathan B Grimm (Grimm et al., 2015; Grimm et al., 2017) and were incorporated into worms by feeding based on (Wu et al., 2019) with the following modifications. L4 worms were washed and collected in small eppendorf tubes with 200 µl M9, concentrated OP50, and 2.5 µM HaloTag dye. Tubes rotated at RT for about 17 hours, and worms were then placed on fresh OP50 plates for at least 4 hours to reduce the background signal of the unbound HaloTag ligand.

For imaging experiments using the heat shock inducible DPY-27::GFP, worms were grown to young adult stage and heat-shocked for 1 hr at 35°C, recovered at RT for 8 hr (unless otherwise labeled). Worms were settled in M9 at 4°C for 10 min, and 40 µl were transferred to a well depression microscopy slide with the addition of 10 µl of 50 mM levamisole (LGC). After 10 minutes, the worms were transferred onto a 10% agarose pad on a microscope slide and covered with a 1.7 µm objective slide (high precision, no.1.5H, Marienfeld). Excess liquid was removed using a lab tissue (Kimtech precision wipe), and the edges of the objective slide were sealed with a two-component silicone glue (picodent twinsil speed).

### Confocal microscopy and FRAP

Confocal imaging and FRAP were performed on a scanning confocal microscope (Leica SP8) using an HC PL APO 63x 1.3 NA glycerol objective (Leica) and Leica Application Suite X (version 3.5.5.19976). For wGFP, the white light laser was set to 482 nm with 10-15% laser intensity, and the emission detection was set to 488 - 520 nm with a HyD hybrid photodetector and gain of 162%. For JF549, the white light laser was set to 549 nm with 10% laser intensity, and the emission detection was set to 554-651 nm with a HyD detector and gain of 200% and gating between 0.3 - 6.0. For JF635, the white light laser was set to 633 nm with 10% laser intensity, and the emission detection was set to 638-777 nm with a HyD detector and gain of 100% and gating between 0.30 - 6.00.

For FRAP in the intestine nuclei, 20 pre-bleach images were acquired, followed by a point bleach (smallest possible bleach spot) of 700 ms with 100 % laser power and subsequent acquisition of ∼500 recovery images using 10-15% laser power. The scan speed was set to 600 Hz, with bidirectional scanning (phaseX: 29.752) in a frame size of 256 × 256 pixels (Pixel dwell time 0,002425 s). The pinhole was set to 1 AU, and a 7x digital zoom was used to zoom in to single intestine nuclei of young adult worms. The FRAP experimental protocol can be found here: https://dx.doi.org/10.17504/protocols.io.bpkymkxw

### FRAP data analysis

Image analysis of the fluorescence recovery at the bleach point was performed using a custom-written script in MATLAB (MathWorks). First, lateral drift in pre- and post-bleach image stacks was corrected using DFT-based sub-pixel image registration (Guizar-Sicairos et al., 2008). The area of each intestine nucleus was then manually segmented. The bleached region was determined by automated thresholding (Otsu’s Method) of an image of the difference of the mean pre-bleach images and the mean of the first five post-bleach images. Acquisition bleaching was detected in the mean intensity of the whole nucleus region of interest in the post-bleach images. This decrease in intensity was fitted with a monoexponential decay and used to correct the acquisition bleaching during fluorescence recovery. To correct for differences in initial intensity and extent of photobleaching, such that different datasets could be directly compared, each acquisition bleaching corrected curve was then normalized to an initial value of 1 and an immediate post-bleach value of 0. To estimate the fraction of fluorescent proteins that can diffuse into the bleached region during the experiment’s time course (mobile fraction) and the recovery time constant (*τ*), the post bleach recovery was fitted with monoexponential function with nonlinear least-squares-based fitting. The mobile fraction was calculated from the monoexponential fit at each experiment’s last recorded recovery time point. The recovery half-time (t1/2), corresponding to the time required to recover half of the fluorescence maximum, is estimated directly from the data. The mean normalized relative intensity of all repeats for each experimental condition was calculated and plotted for each time point with the standard error of the mean (s.e.m.) using Python. The MATLAB analysis script can be found here: https://github.com/ercanlab/2021_Breimann_et_al

### Intensity distribution analysis

To compare the protein expression and X-enrichment of DPY-27::GFP and DPY-27(EQ)::GFP images were recorded at 3 and 8 hours after a 1-hour heat-shock at 35°C. 2D images were manually segmented for the nuclear region, and pixel intensity values for the GFP tagged proteins were recorded for at least 20 images per condition. To compare the average density of pixel intensities per condition, the pixel intensities were binned to ranges of 20, summed for all images of one condition, and divided by the number of used images using Python. To compare image intensities of endogenous DPY-27::Halo in wild-type and *dpy-21 null* conditions, worms were stained with HaloTag-JF549, as described above, and z-stack images were recorded to capture the complete intestinal nuclei. To compare DPY-27::Halo enrichment at the X chromosome between different conditions, the HaloTag signal was segmented in 3D using autocontex pixel classification in ilastik, resulting in a simple segmentation that assigns the most probable class for each pixel (Berg et al., 2019). Using Fiji (Schindelin et al., 2012), a binary 3D mask was created from the ilastik segmentation using Otsu’s method and used to segment the HaloTag signal. Binned pixel intensities were recorded from both conditions, and density plots were created using Python https://github.com/ercanlab/2021_Breimann_et_al.

### Recombinant protein and peptide binding assay

The DNA encoding for amino acids 351-661 of the DPY-28 protein was amplified from cDNA using the primers DPY 28 351F & DPY-28 660R (Table S2). The cDNA template was prepared from total RNA using SuperScript III (Invitrogen) according to the manufacturer’s protocol. The PCR product was digested with BamHI and EcoRI and cloned into corresponding sites in pGEX-5X-2. The plasmid was transformed to a BL21 codon + *E. coli* strain to be induced with 1 mM IPTG for 3 hours at 25°C and purified using standard GST protein purification using GE Healthcare Glutathione Sepharose 4B based on the manufacturer’s protocol, and the protein amount was quantified using a Bradford assay. The peptides were kindly provided by Brian Strahl (Fig. S3D). Briefly, 60 µl of the magnetic streptavidin beads (Dynabeads M280; Invitrogen) were washed twice with 1 ml recombinant protein binding buffer (rPBB) (50 mM Tris pH 8, 0.3 M NaCl, 0.1% Igepal CA360) and incubated rotating 1 hour with 1 nmol peptide at 4°C. The beads were washed twice with rPBB and incubated with 40 pmol of recombinant protein for 3 hours, rotating at 4°C. The beads were washed 5 min thrice with rPBB and resuspended in 30 µl SDS sample buffer, and 15 µl was run on a 4-12% Bis-Tris MOPS gel (Invitrogen) transferred to a PVDF membrane and was blocked with 1xPBST (0.1%Tween-20) containing 5% dry milk. Bound peptides were visualized using an anti-GST antibody (GE 27-4577-50) 1:2,000, Anti-goat-HRP (Promega V8051) 1:10,000 ECL-Plus (GE), and the Typhoon Scanner.

## Acknowledgments

We thank Barbara Meyer for the *dpy-21(JmjC)* strain, and Brian Strahl, Scott Rothbart for peptides and in-solution peptide binding protocol, and Luke Lavis, Jonathan Grimm for the HaloTag dyes. We thank Lennon Matchett-Oates, Parvathy Manoj, Mandy Terne, and Lara Winterkorn for their contributions to the project in general, and Ibai Irastorza Azcarate and Gesa Loof for help with data visualization. We thank the sequencing and imaging facilities at the NYU Biology and the Center for Genomics and Systems Biology, and the Berlin Institute for Medical Systems Biology (BIMSB). We want to thank the MDC-NYU PhD Exchange program for enabling this collaboration.

