## Supplemental Information for "The H4K20 demethylase DPY-21 regulates the dynamics of condensin DC binding"

### Supplementary information

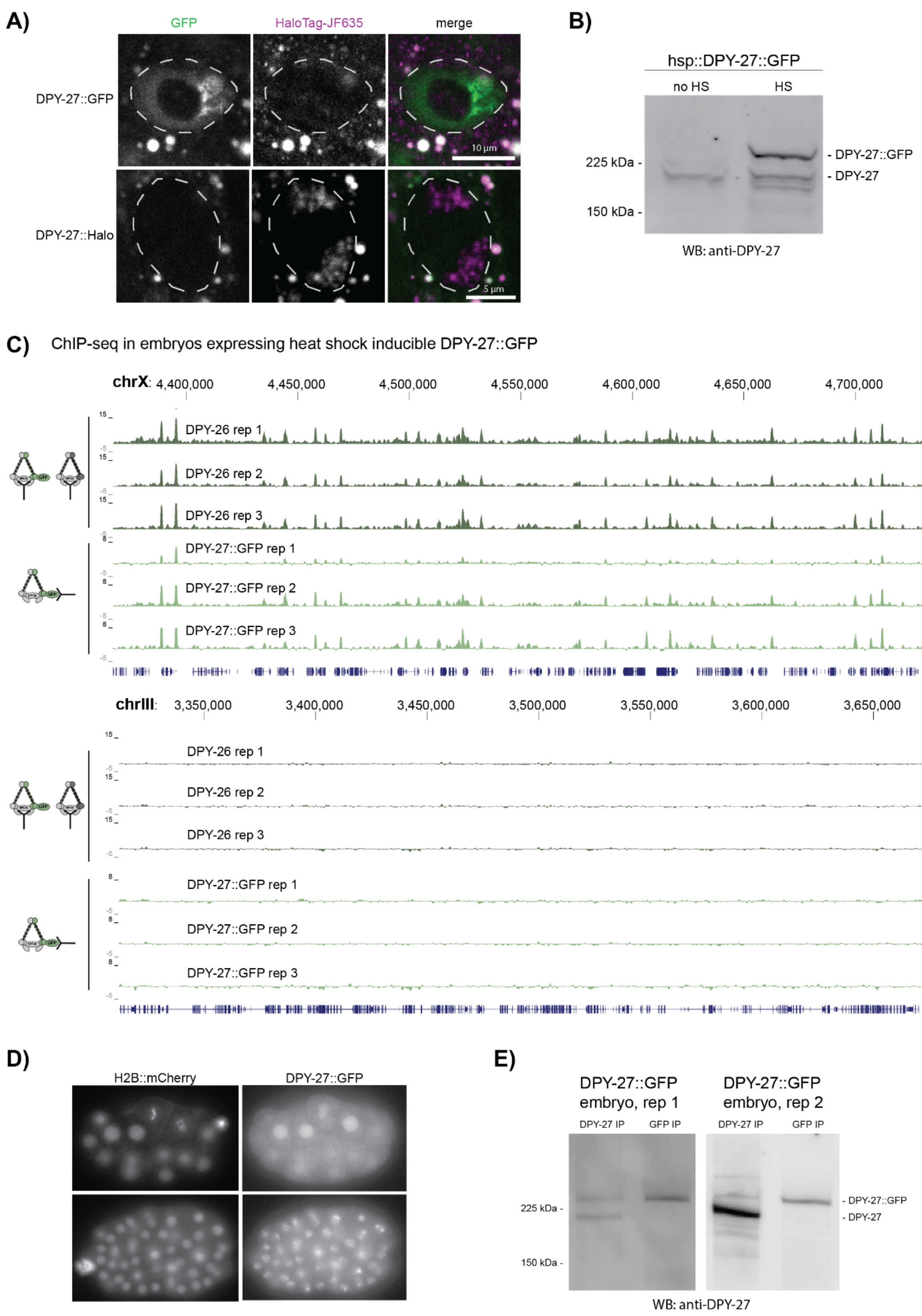

**Figure S1**

**A)** Validation of separable GFP and JF635-Halo signal. Fluorescent images of intestine nuclei after feeding JF635-Halo ligand in homozygous worms expressing heat-shock inducible DPY-27::GFP (upper row) or endogenously Halo-tagged DPY-27 (lower row). Both worm lines were stained with JF635-Halo ligand and heat-shocked. White dotted lines mark nuclei.

**B)** Protein extracts prepared from heat-shocked (HS) and non-heat-shocked (NHS) young adults carrying the *hsp::dpy-27::gfp* transgene were used for western blot. Incubation with DPY-27 antibody shows specific DPY-27::GFP expression upon heat shock.

**C)** Validation of DPY-27::GFP localization specifically to the X chromosomes by ChIP-seq. DPY-27::GFP ChIP-seq analysis replicates using an anti-GFP antibody in embryos. DPY-26 ChIP-seq was used as a positive control in the same extracts.

**D)** X-localization of DPY-27::GFP in embryos is indicated by subnuclear puncta that appears later in embryogenesis when condensin DC localizes specifically to the X chromosomes. H2B::mCherry and DPY-27::GFP signal 6 hours after a heat shock in early (before X localization and late embryos (after DC localization to the X).

**E)** Heat shock expression of DPY-27::GFP was variable in embryos. Two examples are shown where immunoprecipitation of DPY-27 showed different proportions of GFP tagged DPY-27 (top band) compared to endogenous (bottom band). Protein extracts were prepared from embryos isolated from gravid adults that were heat-shocked for 30 min at 35°C and recovered at room temperature for 2 hours.

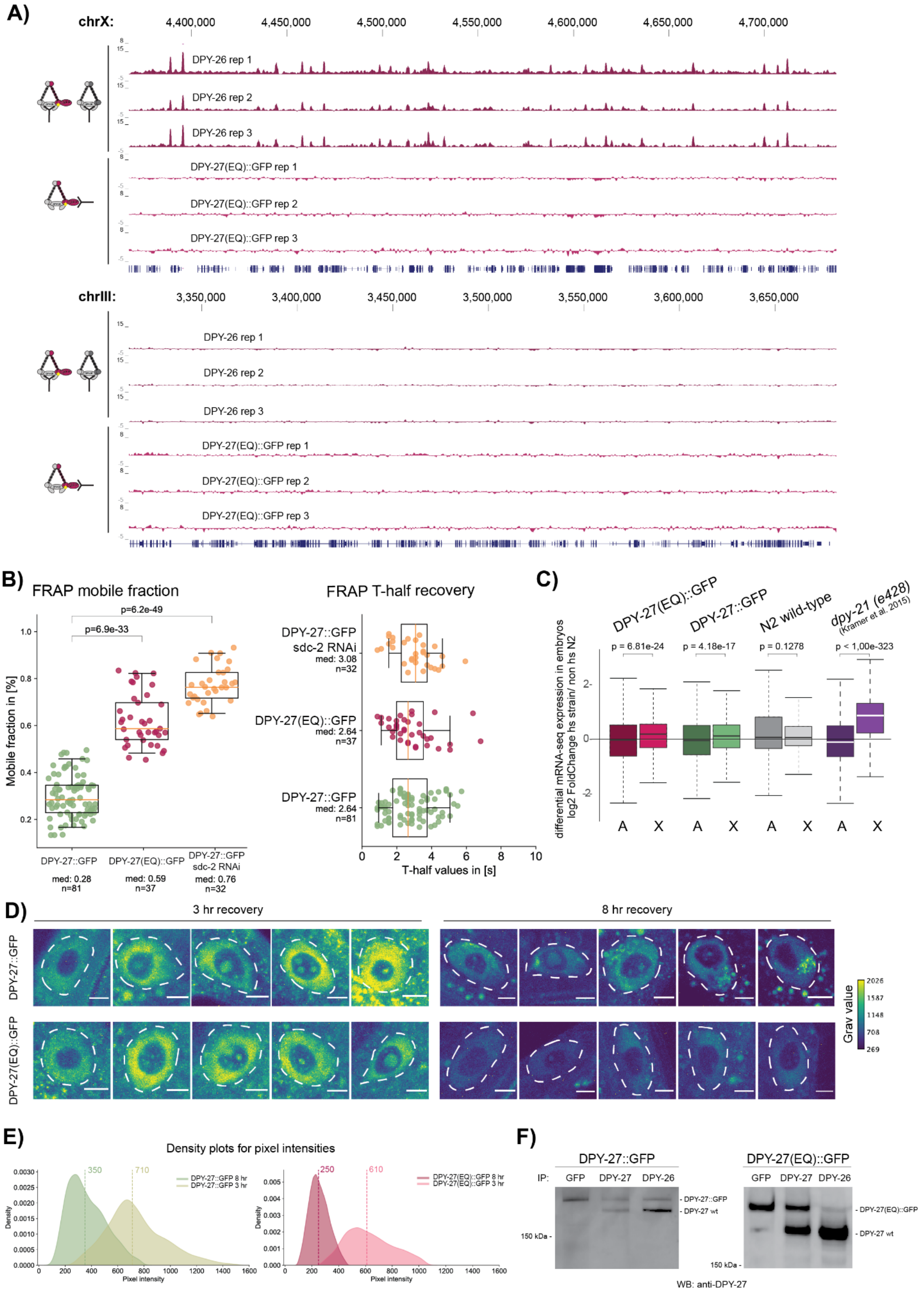
A)
chrX:
DPY-26 rep 1
DPY-26 rep 2
DPY-26 rep 3
DPY-27(EQ)::GFP rep 1
DPY-27(EQ)::GFP rep 2
DPY-27(EQ)::GFP rep 3
chrIII:
B)
FRAP mobile fraction
Mobile fraction in [%]
DPY-27::GFP med: 0.28 n=81
DPY-27(EQ)::GFP med: 0.59 n=37
DPY-27::GFP sdc-2 RNAi med: 0.76 n=32
p=6.2e-49
p=6.9e-33
FRAP T-half recovery
T-half values in [s]
C)
DPY-27(EQ)::GFP
DPY-27::GFP
N2 wild-type
dpy-21 (e428)
(Kramer et al. 2015)
differential mRNA-seq expression in embryos log2 FoldChange hs strain/ non hs N2
p = 6.81e-24
p = 4.18e-17
p = 0.1278
p < 1,00e-323
D)
3 hr recovery
8 hr recovery
DPY-27::GFP
DPY-27(EQ)::GFP
Gray value
E)
Density plots for pixel intensities
Density
Pixel intensity
DPY-27::GFP 8 hr
DPY-27::GFP 3 hr
DPY-27(EQ)::GFP 8 hr
DPY-27(EQ)::GFP 3 hr
F)
DPY-27::GFP
DPY-27(EQ)::GFP
IP: GFP DPY-27 DPY-26
- DPY-27::GFP
- DPY-27 wt
- DPY-27(EQ)::GFP
150 kDa -
WB: anti-DPY-27

**Figure S2**

**A)** ChIP-seq data from replicates corresponding to Figure 2C. Replicates for the wild-type DPY-27::GFP ChIP-seq data can be found in Figure S1C.

**B)** FRAP analysis of mobile fractions (left panel) and T-half recovery time (right panel) corresponding to Figure 2D. P values are from an independent two-sample t-test.

**C)** Log2 fold changes in mRNA-seq between heat-shocked strains and non-heat-shocked wild-type embryos, collected after 30 min heat shock at 35°C followed by 2-hour recovery. Additionally, mRNA-seq log2 fold changes of non-heat-shocked *dpy-21 (e428)* from (Kramer et al., 2015). P values are from a Wilcoxon–Mann–Whitney test.

**D)** Example images of heat-shock expression of wild-type DPY-27::GFP and the ATPase mutant DPY-27(EQ)::GFP after 3 and 8 hours of recovery that were quantified in Supplemental Figure 2E. Images are normalized to the same gray values, and the scale bar corresponds to 5 µm.

**E)** Quantification of the GFP signal's pixel intensities in the nuclei after 3 and 8 hours of recovery from heat shock. The intensities were recorded from at least three biological replicates in adult intestine cells. For wild-type, DPY-27::GFP 21 images were used for the 3-hour intensity curve and 26 for the 8 hours recovery time point. For the intensity curves of DPY-27(EQ)::GFP, 44 images were used for the short time point and 36 images for the long recovery time point. Dotted lines indicate the median value for each distribution.

**F)** Co-immunoprecipitation analysis of condensin DC subunits in embryos. Protein extracts were prepared from embryos that were heat-shocked for 1 hour at 35°C and recovered at room temperature for 2 hours. Immunoprecipitated (IP) DPY-27::GFP and endogenous protein were analyzed by blotting with an anti-DPY-27 antibody. The intensity of the DPY-27::GFP and endogenous protein bands indicate their abundance in each immunoprecipitation.

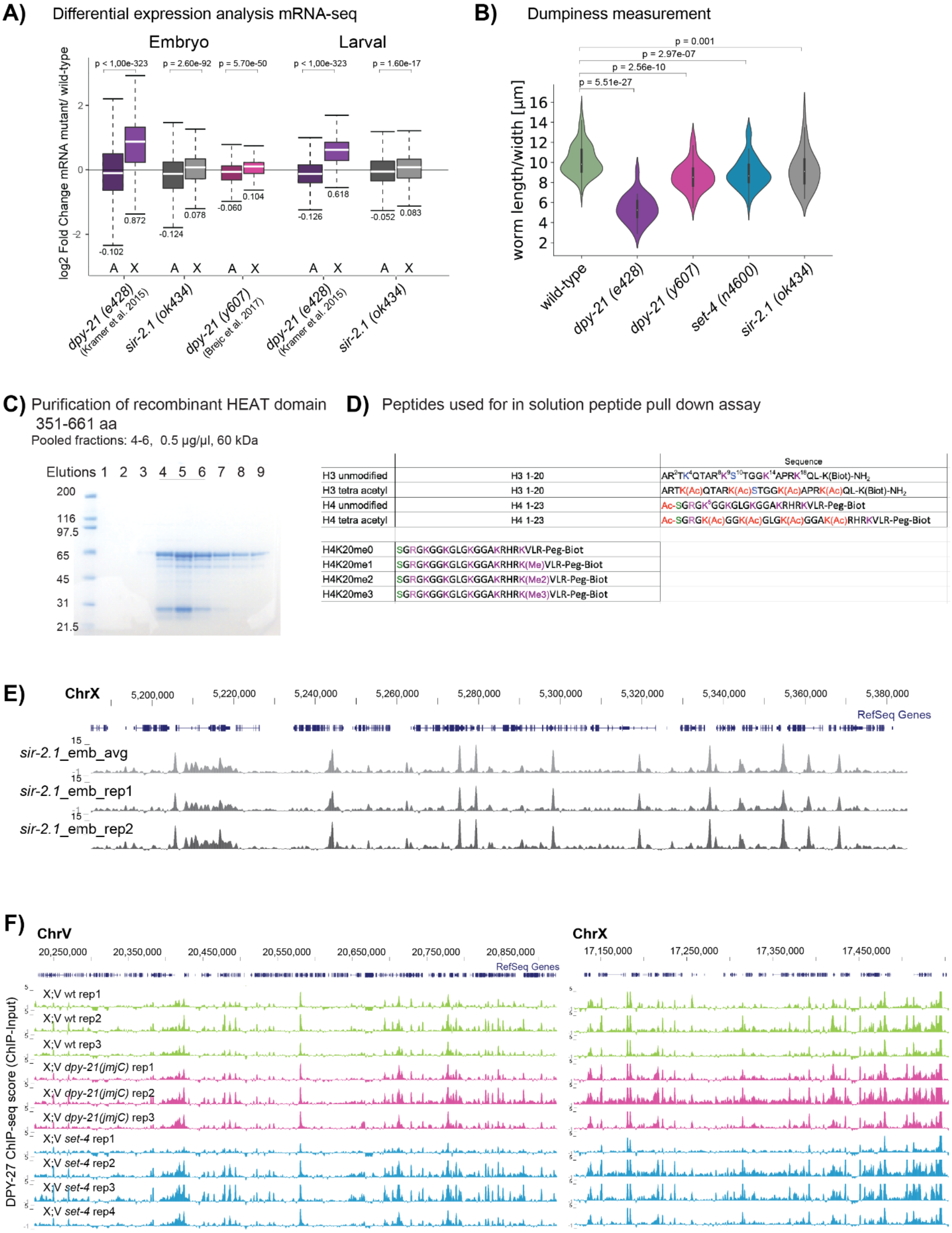

**Figure S3**

**A)** mRNA-seq analysis comparing published *dpy-21 (e428)* from (Kramer et al., 2015), *dpy-21(JmjC)* data from (Brejc et al., 2017), and new data in *sir-2.1* null mutant in embryos (left) and larvae (right). The level of X chromosome derepression compared to autosomes was compared in different mutants using log2 expression ratios compared to wild-type. Significant X chromosome upregulation was tested by a Wilcoxon–Mann–Whitney test. Median values of each group of genes are shown below each boxplot.

**B)** Dumpiness phenotype analysis of wild-type and different mutant worms. The length divided by the width of young adult worms was calculated as a proxy for their dumpiness level from two biological replicates. The following number of worms were used for each condition: wild-type: n= 102; *dpy-21(e428)*: n= 24; *dpy-21(y607)*: n= 67; *set-4 (n4600)*: n= 69; *sir-2.1(ok434)*: n= 51. P values are from an independent two-sample t-test.

**C)** Elutions of GST-DPY-28 HEAT repeat domain recombinant protein, predicted to be ~60 kDa. Fractions 4-6 were pooled for peptide binding assay.

**D)** Sequences and modifications of the N-terminal histone peptides analyzed.

**E)** UCSC genome browser shot of a representative region showing similar DPY-27 ChIP-seq patterns in *sir-2.1* replicates.

**F)** UCSC genome browser shot of replicates corresponding to Figure 3F. Genome browser view of DPY-27 ChIP-seq enrichment on the X chromosomal region of the X;V chromosome in X;V wild-type, *dpy-21(JmjC)* and *set-4* null backgrounds.

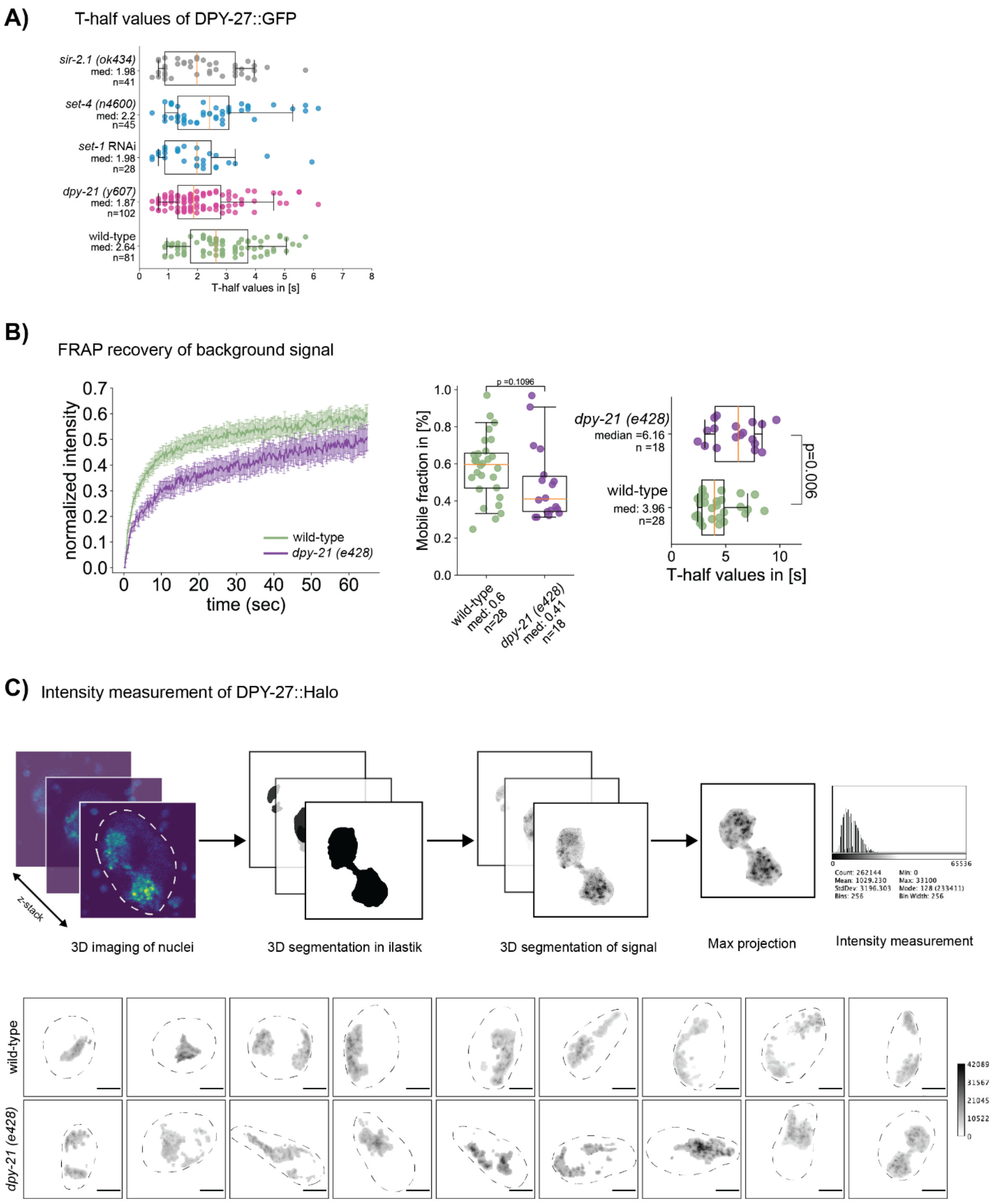

**Figure S4**

**A)** T-half recovery time calculated from individual replicate FRAP recovery curves in Figure 4A. The T-half value for *dpy-21 (e428)* is not included in the plot due to the very low recovery during the experimental time frame.

**B)** Mean FRAP recovery curves for background DPY-27::GFP in wild-type and *dpy-21 (e428)* mutant worms. FRAPs were performed 8 hours after a 1-hour heat shock at 35°C. Unlike Figure 4A, the bleach point was not placed outside the X chromosomal area. Error bars for the bleach curves denote s.e.m. Number of bleached single intestine nuclei (from 2 biological replicates) for each experiment is n = 28 for wild-type and n = 18 for *dpy-21 (e428)*. The middle panel depicts the mobile fractions for the background recovery. The right panel depicts the T-half recovery times for the background recovery. P values are from an independent two-sample t-test.

**C)** Analysis of the fluorescence intensity of endogenously tagged DPY-27::Halo, for wild-type and *dpy-21 (e428)* worms corresponding to Figure 4C. The top row shows the analysis pipeline for the 3D segmentation of the HaloTag-JF549 signal. Z-stacks of intestine nuclei were imaged and segmented in 3D using ilastik (Berg et al., 2019). The resulting mask was used to segment the fluorescent signal in 3D, and from max projections, binned intensities were obtained. The bottom row depicts example images of 3D segmented and max projected nuclei of the wild-type and mutant worms. The scale bar corresponds to 5 µm, and all images are calibrated to the same gray values.

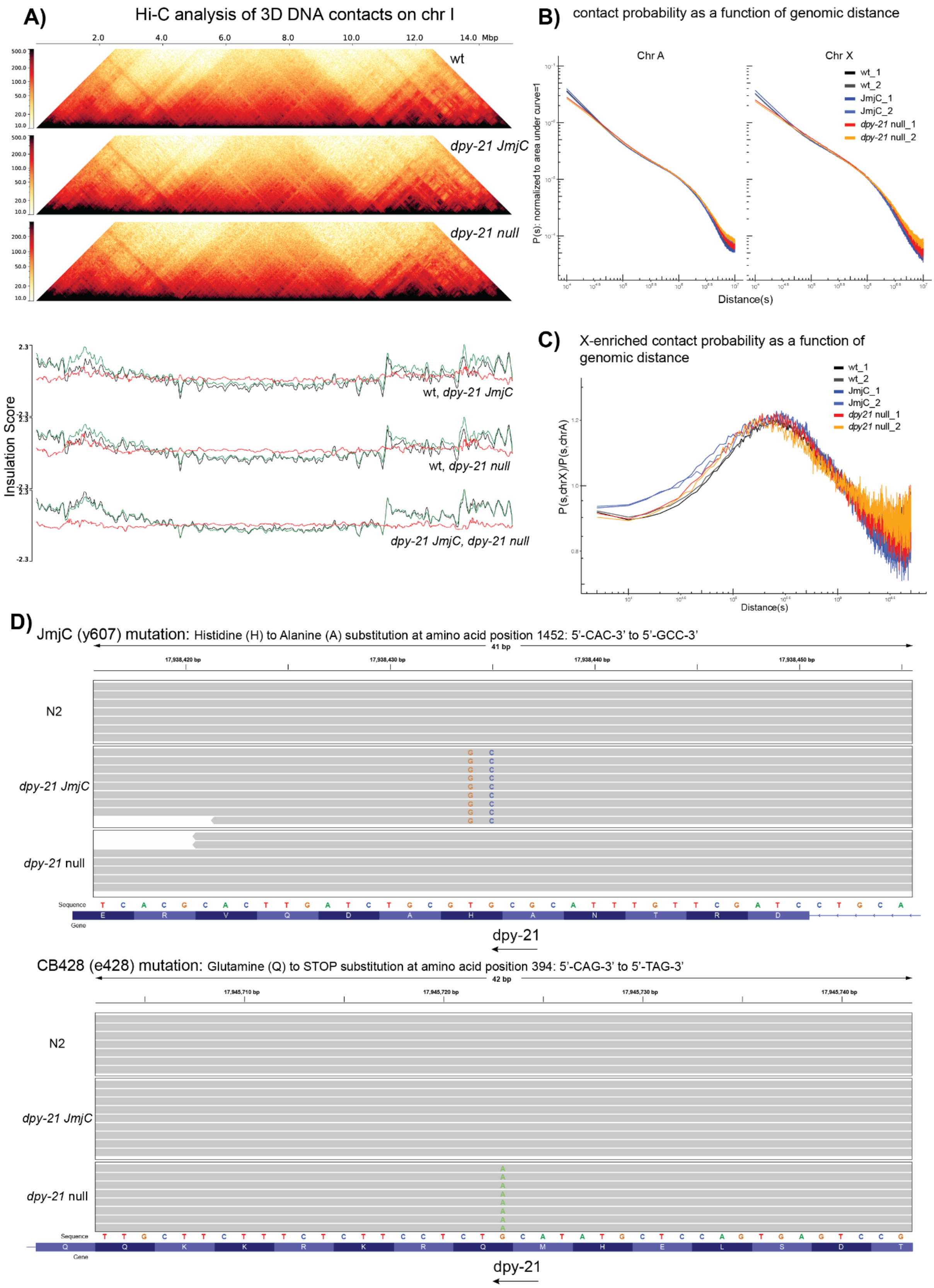

**Figure S5**

**A)** The same plot as Figure 5A for chromosome I.

**B)** Distance decay curve showing the relationship between 5- kb binned genomic separation, s, and average contact probability, P(s) for two biological replicates for each condition.

**C)** The same plot as Figure 5D for each replicate.

**D)** Hi-C reads were used to check for the validity of the strains. The reads were mapped to the ce10 genome. IGV snapshot of mapped reads sorted by mapping quality. Going top to bottom, the three samples correspond to the following genotypes: N2, *dpy-21 JmjC*, and *dpy-21 null*.

**Table S1. List of the C. elegans strains used in this study**

| *strain name* | *strain genotype* | *RRID of strain* | *short description* |
|---|---|---|---|
| N2 | wild type | RRID:WB_STRAIN:N2 | wild type laboratory strain |
| CB428 | dpy-21(e428) V | RRID:WB_STRAIN:CB428 | dpy-21 null |
| YPT47 | X;V | RRID:WB_STRAIN:YPT47 | X;V fusion in wild type background |
| MT14911 | set-4 (n4600) II | RRID:WB_STRAIN:MT14911 | set-4 null |
| ERC47 | ersSi12[hsp16-41::dpy-27::GFP::3xFlag, unc-119(+)] II; unc-119(ed3) III | | promoter_hsp::dpy-27::GFP in mossci Chr II site |
| ERC55 | ersSi21[hsp16-41::dpy-27[EQ-TR]::GFP:: 3xFlag, unc-119(+)] II; unc-119(ed3) III | | promoter_hsp::dpy-27 EQ TR mutation::GFP in mossci Chr II site |
| ERC57 | set-4 (n4600) II; X;V (ypT47) | | X;V fusion in set-4 null background |
| VC199 | sir-2.1(ok434) IV | RRID:WB_STRAIN:VC199 | sir-2.1 null |
| ERC76 | ers53[dpy-27::halo] III | | dpy-27::halo endogenous location fully complementing function with tag and 11 aa deletion |
| ERC81 | dpy-21(y607)V; X;V (ypT47) | | X;V fusion in dpy-21(*JmjC*) background (as in Brejc et al. 2017, introduced into X;V endogenously using CRISPR/Cas9) |
| TY5686 | dpy-21(y607) | | dpy-21*(JmjC*) catalytic mutant (Brejc et al. 2017) |
| RW10993 | unc-119(ed3) III; itIs37 IV; stIs10116; wgIs94. | | H2B-mCherry |
| ***strains derived for FRAP analysis by crossing*** | | | |
| SPL7 | RW10993 X ERC55 | | |
| SPL8 | RW10993 X ERC47 | | |
| SPL13 | SPL8 X TY5686 (*dpy-21* y607 *JmjC* (codon 1454 changed to GCT from GAT) ) | | |

| | |
|---|---|
| SPL14 | SPL8 X MT14911 (*set-4* null) |
| SPL15 | SPL8 X CB428 (*dpy-21* null) |
| SPL16 | SPL8 X VC199 |
| SPL17 | ERC47 X ERC76 |
| SPL18 | RW10993 X ERC76 |
| SPL19 | CB428 X ERC76 |

#### Table S2. List of the primers used in this study

| Target / purpose | F primer | R primer | Forward Sequence | Reverse Sequence | Product Size (bp) |
|---|---|---|---|---|---|
| amplify DPY-28 to be cloned into pGEX5X-2 | DPY 28 351F | DPY-28 660R | atgcGGATCCGAACGAGCCGAAAAGCCA | atgcGAATTCTCAATCGTCCATTTGGGTTAG | |
| hsp promoter amplification from pCM1.57 with overlapping 15 bp to XhoI cut pCFJ151 for in fusion cloning, reverse primer is complementary to start of DPY-37 3' UTR | SE123F | SE123R | GCAGGAATTCCTCGActgcaggtcgactctaga | atatattacttcaatatttttctaccggtacc | ~550 bp |
| dpy-27 3'UTR amplification with overlap to pCJF151 on the right side | SE124F | SE124R | attgaagtaatatattttaaac | CACCGTACGTCTCGAttaggaaattatttttgat | ~400 bp |
| amplify DPY-27 on the left overlapping with pCFJ151 with hsp promoter | SE135F | SE135R | gtattggtaccggtagaaaaaatATGCAGCCGTTTAAAAGACG | ATGTCTGCTTCTTCGCACAC | ~5.8 kb |
| amplify GFP3xflag on the left overlapping with DPY-27 and on the right with PCFJ151 dpy27 3' UTR | SE136F | SE136R | CGAAGAAGCAGACATATGAGTAAAGGAGAAGAACTT | gtttaaaatatattacttcaatTCACTTGTCATCGTCATCC | ~1 kb |
| DPY-27 clone sequencing verification | SE127 | | gatgacgacaagggcagc | | |
| DPY-27 clone sequencing verification | SE128 | | cgggcctgagatccacac | | |
| DPY-27 clone sequencing verification | SE129 | | GCAAAGGATGAAGTTCGG | | |
| DPY-27 clone sequencing verification | SE130 | | TGAACTGAAAGAGGCTGG | | |
| DPY-27 clone sequencing verification | SE131 | | ATCACGCACTGGAAGCTC | | |
| DPY-27 clone sequencing verification | SE132 | | AATTGCAAACTTCAACGg | | |
| DPY-27 clone sequencing verification | SE133 | | AAGATGTTGACAAGTTCC | | |
| DPY-27 clone sequencing verification | SE134 | | TCGAGGAGGAGATCAAAC | | |
| Q5 mutagenesis of DPY-27 sequence for E-Q mutation | SE101F | SE101R | cAgATCGATGCGGCACTGGAC | ATCCATCACGTAGAGGGGTG | |
| crRNA from IDT to cut at 3' end of *dpy-27* | LS37 | | ACACGGCGTTGAACGACAAT | | |
| amplify halo from pLS19 with homology arms to tag DPY-27 | AM29F | AM29R | CATCTCCACCACCAATCGTCGTCCAACG | ttttcaaaatttagtttaaaatatattacttcaatTTATCCGGAG | |

| | | | | |
|---|---|---|---|---|
| C terminus. 5' SP9 modified primers | | | TCGCGTGCGAAGAAGCAGACATgggggaggaggatcgGAAATCGGTACAGGCTTTCC | ATCTCGAGGGTGG |
| primers to detect halo insertion at 3' end of *dpy-27* | LS40F | LS40R | TGGACAGTACGTGATGCAAAG | CGATGAGCCAGTAAGAAGACG |
| crRNA from IDT for JmjC dpy-21 H1452A | BR16_sgRNA | | TTTCGACCTGAAATTTCACG | |
| oligo repair template for JmjC dpy-21 mutant | BR16_oligo | | TGGATCATCTTCAGTTGATTCACGCACTTGATCTGCGGCCGCATTTGTTCGATCCTGCACAAAAATAGTTGAAATTTGAGTTTTTTGTAAATTTTAACAGTTTTTCAATAGAAAATTCGTATTCGTCGTGAAATTTCAGGTCGAAATGGGTTTTTTTTCGAAAACATTTGTGGTTGAAAAAGTGGCTCAGTCGGTAAGAT | |
| Amplify region of dpy-21 JmjC mutantion (product size 514bp. Not1 digestion products: 216bp - 298bp) | BR17F | BR17R | AACTATTGACCACACCCGGG | GGCGGTTCGTAGAGATCCAT |

#### Table S3. List of the antibodies used in this study

| Target | Antibody | Antibody information | Antigen | RRID of antibody | Reference |
|---|---|---|---|---|---|
| DPY-27 | JL00001 | Rabbit polyclonal | 1-409 aa | Covance Research Products Inc Cat# JL00001_DPY27, RRID:AB_2616039 | Ercan et al 2007 Nature Genetics |
| DPY-26 | JL00003 | Rabbit polyclonal | 740-1262 aa | Covance Research Products | Ercan et al 2009 Current Biology |
| MIX-1 | JL00004 | Rabbit polyclonal | 837–1244 aa | Covance Research Products | Ercan et al 2009 Current Biology |
| GFP | ab290 | Rabbit polyclonal | Recombinant full-length protein corresponding to GFP. Green fluorescent protein (GFP) from Aequorea victoria. | Abcam | -- |

**Table S4. Information for RNA-seq data used in this study**

| **RNA-seq data from this study** | | | | | | |
|---|---|---|---|---|---|---|
| ***GEO accession number*** | ***Sequencing ID*** | ***Description*** | ***Strain*** | ***Stage*** | ***Mapped reads*** | ***Technical Reps*** |
| GSM5075626 | SEA51 | VC199_emb_rep1A | VC199 | mixed embryos | 13,342,597 | SEA58 tech rep |
| GSM5075626 | SEA58 | VC199_emb_rep1B | VC199 | mixed embryos | 21,079,971 | SEA51 tech rep |
| GSM5075627 | SEA70 | VC199_emb_rep2 | VC199 | mixed embryos | 23,574,464 | |
| GSM5075628 | SEA77 | VC199_emb_rep3 | VC199 | mixed embryos | 17,555,806 | |
| GSM5075629 | SEA86 | VC199_emb_rep4 | VC199 | mixed embryos | 20,187,638 | |
| GSM5075630 | MK11 | VC199_L2L3_Rep1A | VC199 | L2-L3 | 15,784,655 | MK25 tech rep |
| GSM5075630 | MK27 | VC199_L2L3_Rep1B | VC199 | L2-L3 | 16,716,809 | MK19 tech rep |
| GSM5075631 | MK19 | VC199_L2L3_Rep2 | VC199 | L2-L3 | 10,497,663 | |
| GSM5075632 | MK46 | VC199_L2L3_Rep3 | VC199 | L2-L3 | 21,498,117 | |
| GSM5075633 | MK60 | VC199_L2L3_Rep4 | VC199 | L2-L3 | 24,037,603 | |
| GSM5075634 | SEA224 | KB01_emb_RNA_rep1 | KB01 | mixed embryos | 8,220,567 | |
| GSM5075635 | SEA225 | KB01_emb_RNA_rep2 | KB01 | mixed embryos | 9,726,617 | |
| GSM5075636 | LAS41 | KB01_emb_RNA_rep3 | KB01 | mixed embryos | 24,873,910 | |
| GSM5075637 | SEA221 | MK14_emb_RNA_rep1 | MK14 | mixed embryos | 9,012,229 | |
| GSM5075638 | SEA222 | MK14_emb_RNA_rep2 | MK14 | mixed embryos | 10,363,195 | |
| GSM5075639 | SEA223 | MK14_emb_RNA_rep3 | MK14 | mixed embryos | 7,729,455 | |
| GSM5075640 | LAS47 | LS_ext534_emb_RNA_N2_HS_rep1 | N2, heat shock | mixed embryos | 25,478,937 | |
| GSM5075641 | LAS48 | LS_ext536_emb_RNA_N2_HS_rep2 | N2, heat shock | mixed embryos | 18,412,383 | |
| GSM5075642 | LAS49 | LS_ext549_emb_RNA_N2_HS_rep3 | N2, heat shock | mixed embryos | 24,768,721 | |
| **Published RNA-seq data used in this study** | | | | | | |
| ***GEO accession number*** | ***Strain*** | ***Description*** | ***Stage*** | ***Reference*** | | |
| GSE67650 | N2 | N2 | mixed embryo | Kramer et al PLoS Gen 2015 | | |
| GSE67650 | CB428 | dpy-21 null | mixed embryo | Kramer et al PLoS Gen | | |

| | | | | |
|---|---|---|---|---|
| | | | | 2015 |
| GSE67650 | N2 | N2 | L3 | Kramer et al PLoS Gen 2015 |
| GSE67650 | CB428 | dpy-21 null | L3 | Kramer et al PLoS Gen 2015 |
| GSE84581 | N2 | N2 | mixed embryo | Brejc et al Cell 2017 |
| GSE84581 | TY5686 | dpy-21 (y607) | mixed embryo | Brejc et al Cell 2017 |

**Table S5. Sanger sequencing results for ERC76**

| ***Sequencing results for ERC76, Halo CRISPR tagging dpy-27, revealed insertion of unknown sequence (grey) before the tag sequence, which does not affect dpy-27 function*** | |
|---|---|
| ATTGATCGAAGAAGCAACTCCATCTCCACCACCAATCCACTCCTCGTCGAAGGTCGAAATCGGTAC<br>AGGCTTTCCATTCGACCCCCATTATGTGGAGGTCCTCGGAGAGCGTATGCACTACGTCGACGTCGG<br>ACCACGTGACGGAACCCCAGTCCTCTTCCTCCACGGAAACCCAACCTCCTCCTACGTCTGGCGTA<br>ACATCATCCCACACGTCGCCCCAACCCACCGTTGCATCGCCCCAGACCTCATCGGAATGGGAAAGT<br>CCGACAAGCCAGACCTCGGATACTTCTTCGACGACCACGTCCGTTTCATGGACGCCTTCATCGAGG<br>CCCTCGGACTCGAGGAGGTCGTCCTCGTCATCCACGACTGGGGATCCGCCCTCGGATTCCACTGGG<br>CCAAGCGTAACCCAGAGCGTGTCAAGGTAAGTTTAAACATATATATACTAACTAACCCTGATTATTT<br>AAATTTTCAGGGAATCGCCTTCATGGAGTTCATCCGTCCAATCCCAACCTGGGACGAGTGGCCAGA<br>GTTCGCCCGTGAGACCTTCCAAGCCTTCCGTACCACCGACGTCGGACGTAAGCTCATCATCGACCA<br>AAACGTCTTCATCGAGGGAACCCTCCCAATGGGAGTCGTCCGTCCACTCACCGAGGTCGAGATGG<br>ACCACTACCGTGAGCCATTCCTCAACCCAGTCGACCGTGAGCCACTCTGGCGTTTCCCAAACGAG<br>CTCCCAATCGCCGGAGAGCCAGCCAACATCGTCGCCCTCGTCGAGGAGTACATGGACTGGCTCCA<br>CCAATCCCCAGTCCCAAAGCTCCTCTTCTGGGGAACCCCAGGAGTCCTCATCCCACCAGCCGAGG<br>CCGCCCGTCTCGCCAAGTCCCTCCCAAACTGCAAGGTAAGTTTAAACAGTTCGGTACTAACTAACC<br>ATACATATTTAAATTTTCAGGCCGTCGACATCGGACCAGGACTCAACCTCCTCCAAGAGGACAACC<br>CAGACCTCATCGGATCCGAGATCGCCCGTTGGCTCTCCACCCTCGAGATCTCCGGATAAattgaagtaata<br>tattttaaactaaattttgaaaaaaaaaagaaaactttgtgaaaaatccaaaaaatgagaccaactttcttt | |
| dpy-27 | [blue] |
| insertion | [grey] |
| halo | [green] |
| stop codon | [red] |

**Table S6. Sanger sequencing results for ERC81**

| ***Sequencing results showing CAC to GCC change to generate the X;V, dpy-21(JmjC y607)*** |
|---|

NNNNNNNNNNNNNNNNNNNNTGGTAGANGACCATCTTACCGACTGAGCCACTTTTTCAACCACA
AATGTTTTCGAAAAAAAACCCATTTCGACCTGAAATTTCACGACGAATACGAATTTTCTATTGAAA
AACTGTTAAAATTTACAAAAAACTCAAATTTCAACTATTTTTGTGCAGGATCGAACAAATGCGGCC
GCAGATCAAGTGCGTGAATCAACTGAAGATGATCCAACAACAACAACAACAACTACAACGACTAC
AAGTTCTTCTTCTTCTTCTTCAAAATCGAAAAAATCGGCGAAATCCGATCCGACATTTGTTAAATCA
ACGGCTGCTGTGGGTGTCCTACAGGGTATCAGGAATCCTGATGCAAATGACGATGATGAATATTATG
AGGATGAACGAAAAGCTGTTAAAGAAGTTATTGTATTTGATGCACATGATTTGCATAAAGTTGCACA
TCATCTTGCAATGGATCTCNNNAAANCCGCCA

| |
|---|
| crRNA |
| disrupted PAM site |
| changed codon |

**Table S7. Information for ChIP seq data used in this study**

**Table S8. Information for Hi-C data used in this study**

**Table S9. Readme for DEseq output**

| | |
|---|---|
| **Gene annotations are from WS220 (UCSC genome version ce10).** | |
| **Transcripts Per Kilobase Million (TPM) tab** | |
| TPM: | |
| sample_sums<-apply(data.matrix(fpkm[,-c(1:2)]),2,sum) | |
| tpm<-t(t(fpkm[,-c(1:2)])/sample_sums)*10^6 | |
| **DEseqOutput Tabs** | |
| column name: | |
| gene | name of gene |
| wbid | wormbase ID of the gene |
| baseMean | average of normalized count values |
| log2FoldChange | log2 fold change effect size estimate |
| lfcSE | standard error estimate for the log2 fold change values |
| stat | Wald statistic |
| pvalue | Wald test p-value |
| padj | Benjamini-Hochberg adjust p-value |

**Table S10. TPM replicates**

**Table S11. DEseq Output VC199/N2 embryo**

**Table S12. DEseq Output VC199/N2 L3**

**Table S13. DEseq Output MK14/N2 embryo HS**

**Table S14. DEseq Output KB01/N2 embryo HS**
